# Mechanical confinement drives monocyte-to-macrophage differentiation

**DOI:** 10.64898/2026.03.31.715742

**Authors:** Wei Liu, Xue-Zhu Chen, Hao Zhang, Xue Bai, Yu-Ting Du, Yu-Xin Ji, Run-Yuan Mao, Ya-Jun Wang, Mengyao Sheng, Hai Gao, Fei Xavier Chen, Guangyin Jing, Xinxin Huang, Zhengjun Chen, Yan-Jun Liu

## Abstract

Cells in vivo experience mechanically diverse microenvironments in which physical confinement is a pervasive but poorly understood regulator of their behavior and fate. Whether and how mechanical confinement governs immune cell differentiation remains unknown. Here, we reveal that a mechanical cue — long-term confinement is sufficient to drive monocyte-to-macrophage differentiation through a mechanoepigenetic pathway. In vivo, differentiating monocytes exhibited flattened nuclei in the liver capsule, indicative of confinement by surrounding stromal and parenchymal structures. Using a custom cell confiner to recapitulate this confined niche, we found that confinement induces macrophage-like protrusive architectures, enhances motility, and upregulates macrophage-associated genes in RAW264.7 and THP-1 monocyte-lineage cells. Notably, extending this paradigm to primary murine bone-marrow, human umbilical-cord, and tissue-derived hepatic-associated monocytes yielded similar outcomes, thus enhancing phagocytic capacity, directly demonstrating that mechanical confinement can program monocytes into macrophages. Mechanistically, we found that confinement activates KDM6B, leading to H3K27me3 demethylation, which derepresses macrophage-specific transcriptional programs. Pharmacological inhibition of KDM6B with GSK-J4 restored H3K27me3 and blocked macrophage differentiation both in vitro and in vivo. These findings define a KDM6B–H3K27me3 axis that links nuclear mechanics to transcriptional reprogramming, positioning mechanical confinement as a “super-enhancer–like” cue for engineer macrophage function in therapeutic and bioengineering contexts.

## Introduction

Living tissues present densely packed, spatially restricted niches in which cells continually negotiate physical interactions with the extracellular matrix and neighboring cells^1–7^. Among the diverse mechanical cues that influence cell fate and function, mechanical confinement, defined as the restriction of cell shape and volume by surrounding structure, represents a ubiquitous yet underappreciated feature of most tissues^3^. Confinement naturally arises in dense interstitial spaces, fibrotic regions, and immune or tumor microenvironments, where cells must deform to move through limited spaces^3,8–10^. Such spatial restriction not only alters cellular behaviors, including migration and division^11–14^, but also imposes profound effects on cell identity—affecting cell fate decisions, aging processes, and reprogramming potential^15–20^. Understanding how cells sense and adapt to confinement is therefore crucial for deciphering how mechanical cues regulate cell differentiation, immune activation, and tissue homeostasis.

Immune cells are particularly susceptible to confinement as they migrate, surveil, and differentiate within densely packed tissues^8,21^. Histological analyses reveal that interstitial and vascular spaces in vivo range from submicron dimensions (<1 µm) to tens of micrometers (∼70 µm), imposing substantial physical confinement within tissues ^5,9,10,22,23^. During inflammation, infection, or tumor progression, monocytes and macrophages frequently traverse narrow interstitial gaps, infiltrate collagen-rich matrices, and squeeze through endothelial or epithelial barriers^24–26^. Such confined microenvironments impose persistent physical stress, yet how these forces influence immune cell fate and function remains largely unknown.

Recent studies have begun to reveal how immune cells sense and adapt to confinement. Dendritic cells migrating through restricted spaces detect compression-induced deformation via the ARP2/3–cPLA2–NF-κB pathway, which regulates CCR7 expression and lymph-node homing^8^. Likewise, mechanical confinement at endothelial junctions activate the mechanosensor Piezo1 in neutrophils, triggering Ca²⁺ spikes that are essential for migration into Pseudomonas-infected lungs^27^. These findings highlight mechanical confinement as a potent immunoregulatory signal. Converging evidence further shows that confinement-driven nuclear deformation can initiate epigenetic remodeling. Transient nuclear compression alters H3K9me3 levels and promotes fibroblast-neuron reprogramming^15^; cyclic stretching-induced nuclear deformation triggers a calcium-dependent loss of H3K9me3-marked heterochromatin, enabling chromatin softening and mechanically adaptation^28^; and in mesenchymal stem cells migrating under confinement, pronounced nuclear deformation enhances H3K27 acetylation and drives osteogenic differentiation^16^. Similarly, in sliding hydrogel systems that mimic local confinement, nuclear envelope stretching activates cPLA2, remodels H3K9me3/H3K27me3, and promotes chondrogenic differentiation^29^. Collectively, these studies suggest that confinement-mediated nuclear deformation interface with chromatin remodeling to regulate cell identity. Despite these advances, how mechanical confinement influences immune cell differentiation—particularly monocyte-to-macrophage transition—remains largely unexplored. Although biochemical cues such as cytokines and growth factors are well-established drivers of macrophage development, whether spatial confinement provides an instructive mechanical signal in this process is unknown. Elucidating how immune cells response to spatial restriction is therefore essential for understanding their adaptation to tissue environments and for uncovering new principles of mechanoregulation in immunity.

Here, we set out to determine whether mechanical confinement influences monocyte-to-macrophage differentiation. In vivo, differentiating monocytes displayed markedly elongated and flattened nuclei, indicating sustained nuclear deformation imposed by the dense collagen network and surrounding tissue cells. Using a vertical confinement system that recapitulates tissue-level spatial restriction, we find that prolonged confinement drives RAW264.7 and THP-1 cells toward macrophage-like states, marked by complex protrusive morphologies, enhanced migration, and activation of macrophage transcriptional programs. This response is conserved in primary monocytes from mouse bone marrow, human umbilical cord blood, and liver tissue, which acquire macrophage markers (F4/80, CD163), upregulate key lineage regulators (*Adgre1, Itgam, Mafb*), and display increased phagocytic activity. Mechanistically, confinement-induced nuclear deformation promotes KDM6B-dependent removal of H3K27me3, enabling activation of macrophage-specific gene expression. Genetic or pharmacological disruption of KDM6B blocks H3K27me3 erasure, prevents confinement-induced macrophage differentiation in vitro, and impairs macrophage replenishment in vivo. Together, these findings identify a mechano-epigenetic pathway by which spatial confinement instructs monocyte fate, establishing mechanical confinement as a physiologically essential and engineering-compatible cue for controlling macrophage identity.

## 2 Results

### 2.1 Long-term confinement induces phenotypic transition in monocyte-lineage cell lines

In living tissues, immune cells experience pervasive physical confinement imposed by tissue microarchitecture, which mechanically restricts cellular and nuclear deformation and thereby shapes immune cell behaviors^2,8,27^. The liver capsule comprises a compact collagenous layer enriched with fibroblasts and stromal cells, forming the spatially restrictive interface that encapsulates the hepatic parenchyma^30,31^. This specialized architecture establishes a physiologically confinement microenvironment that imposes sustained mechanical constraints on both infiltrating and resident immune cells^9,32,33^. To examine how immune cells respond to mechanical confinement in vivo, we used two-photon microscopy to visualize *Cx3cr1*-GFP⁺ liver capsular macrophages (LCMs) — a population marked by the chemokine receptor Cx3cr1^32^, which labels circulating monocytes and monocyte-derived macrophages — together with second harmonic generation (SHG) signals from collagen fibers (Fig. 1a). LCMs resided within a densely woven collagen mesh and exhibited a distinctive, octopus-like morphology (Fig. 1a). These cells originate from circulating monocytes that traverse narrow vascular barriers and the compact hepatic interstitium to reach the capsule, thereby encountering substantial mechanical confinement^32,34–36^.

**Figure 1.**
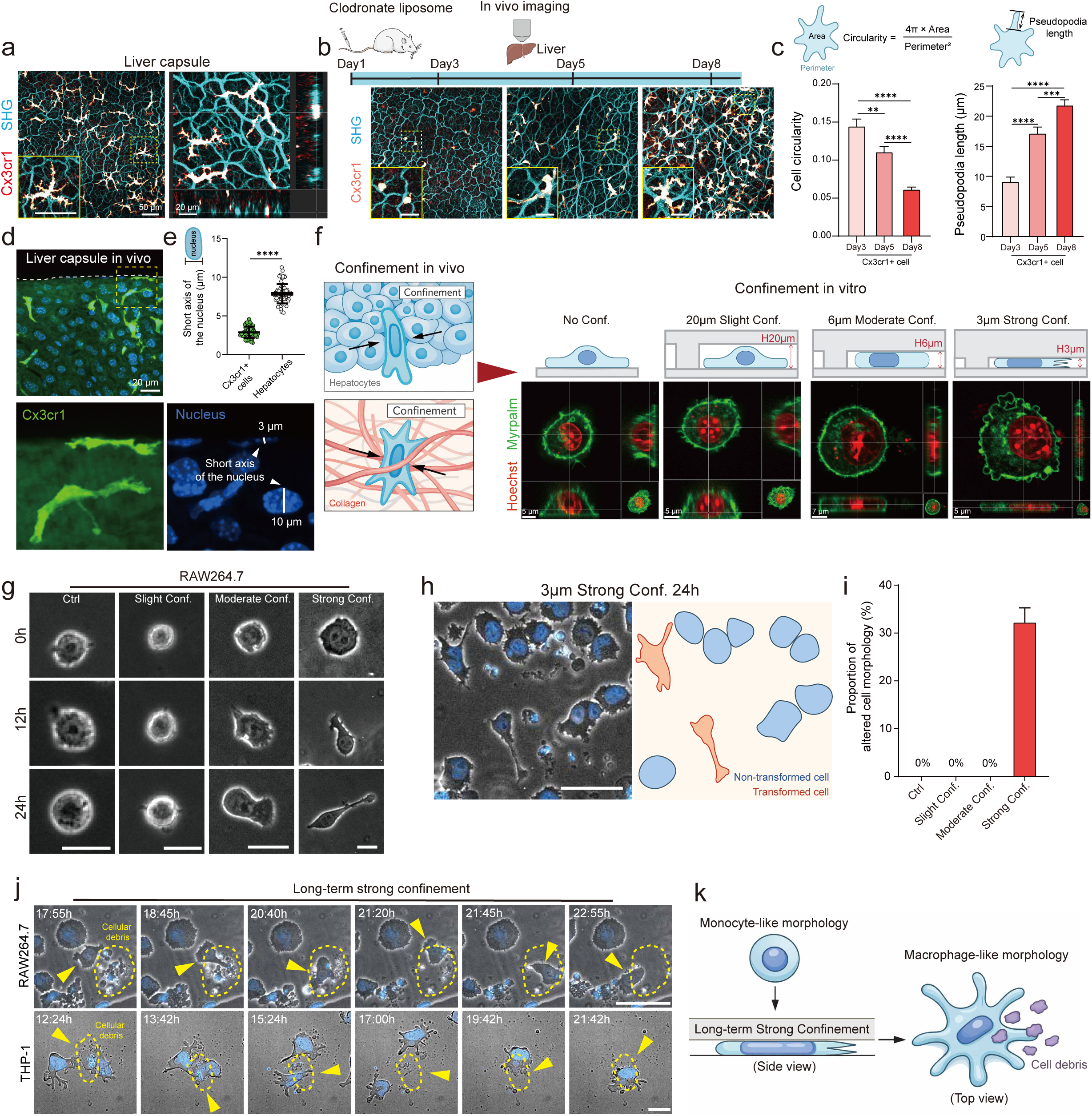
Long-term strong confinement induces phenotypic transition in monocyte-lineage cell lines. a. *In vivo* two-photon intravital imaging of liver capsular macrophages (LCMs) expressing *Cx3cr1* (red–hot). Collagen fibers delineating the dense capsular stromal architecture are visualized by second harmonic generation (SHG) signals (cyan). b. *In vivo* two-photon imaging of *Cx3cr1*⁺ LCMs following clodronate liposome treatment at day 3, day 5, and day 8, with SHG signals (cyan) marking the collagen-rich capsular microenvironment. Scale bar, 20 μm. c. Quantification of cell circularity and pseudopodia length of *Cx3cr1*⁺ LCMs at day 3, day 5, and day 8 after clodronate liposome treatment. Data are shown as mean ± s.e.m. (*N* = 3 independent experiments; *n* = 56 cells for day 3, *n* = 60 cells for day 5, *n* = 60 cells for day 8). Statistical significance was determined by a two-tailed unpaired Student’s *t*-test; **P < 0.01, ***P < 0.001, ****P < 0.0001. d. Immunofluorescence staining of mouse liver tissue sections showing *Cx3cr1*⁺ cells (green) within the hepatic surface region. DAPI labels nuclei (blue). The dashed line marks the boundary of the liver capsule. e. Quantification of the minor nuclear axis (short-axis length) of *Cx3cr1*⁺ cells and neighboring hepatocytes within the liver parenchyma. Data are shown as mean ± s.e.m. (*N* = 3 independent experiments; *n* = 64 *Cx3cr1*⁺ cells, *n* = 74 hepatocytes). Statistical significance was determined by a two-tailed unpaired Student’s *t*-test; ****P < 0.0001. f. Schematic illustration of the confined microenvironment within the liver parenchyma *in vivo* (left) and the in vitro confinement system, showing confinement heights corresponding to 20 μm (Slight Confinement; Slight Conf.), 6 μm (Moderate Confinement; Moderate Conf.), and 3 μm (Strong Confinement; Strong Conf.). g. Representative images of RAW264.7 cells after long-term culture under different confinement conditions: control, slight confinement, moderate confinement, and strong confinement. Increasing confinement height restriction induces progressive morphological alterations. Scale bar, 20 μm. h. Representative images of RAW264.7 cells after 24 h strong confinement (3 μm), showing pronounced morphological changes. Cells undergoing confinement-induced phenotypic transition are outlined in red. Scale bar, 50 μm. i. Quantification of the morphological transition rate of RAW264.7 cells under control, slight confinement, moderate confinement, and strong confinement conditions. Data are shown as mean ± s.e.m. (*N* = 9 independent experiments). j. Representative images of RAW264.7 and THP-1 cells following long-term strong confinement, showing confinement-induced morphological transition and phagocytosis of surrounding cellular debris. Scale bar, 50 μm. k. Schematic model illustrating long-term strong confinement–induced morphological transition and enhanced phagocytic activity in monocyte-lineage cell lines.

To follow the in vivo maturation of LCMs, we depleted liver-resident macrophages with clodronate liposomes (CLL) and monitored the stepwise transition from monocytes to macrophages (Fig. 1b). After CLL-mediated clearance, a few *Cx3cr1*⁺ cells reappeared by day 3, embedded within dense collagen matrices but lacking the elaborate morphology of mature LCMs. Over time, the numbers of *Cx3cr1*⁺ cells increased, and the cells gradually acquired the complex, spread-out morphology characteristic of differentiated macrophages (Fig. 1b).

We performed quantitative morphometric analysis and found that cell spreading area, pseudopodia length, and Feret’s diameter increased, whereas circularity and solidity decreased (Fig. 1c and Extended Data Fig. 1a – c), revealing progressive morphological remodeling during the monocyte-to-macrophage transition. Histological staining further showed that *Cx3cr1*⁺ immune cells at the capsule possessed markedly flattened nuclei, indicating pronounced nuclear deformation, whereas neighboring hepatocytes retained spherical nuclei (Fig. 1d). Quantitative morphometry reinforced this observation: the average nuclear minor-axis length of *Cx3cr1*⁺ cells was ∼3 µm, compared with ∼7 µm in hepatocytes (Fig. 1e), confirming this nuclear deformation.

Together, these findings indicate that monocytes migrating from the vasculature into the collagen-rich hepatic capsule experience strong mechanical confinement and nuclear compression during differentiation into macrophages (Fig. 1f left). Therefore, we speculate that such sustained confinement likely acts as a physical cue that orchestrates cellular morphogenesis and may instruct macrophage fate determination.

To dissect how mechanical confinement regulate monocyte-to-macrophage differentiation, we utilized a previously established vertical confinement system (cell confiner) capable of imposing spatial restriction with micrometer-level precision (Fig. 1f right). We confined RAW264.7 cells labeled with Myrpalm-mEmerald and Hoechst under three defined heights: 20 μm (slight confinement), 6 μm (moderate confinement) and 3 μm (strong confinement) (Fig. 1f). The results showed that 20 μm slight confinement had no significant effect on cell morphology. In contrast, 6 μm moderate confinement resulted in the moderate deformation of both the cell body and nucleus, whereas 3 μm strong confinement caused pronounced deformation of both compartments (Fig. 1f).

Typically, RAW264.7 cells require stimulation with LPS or IL-4 to undergo maturation, transitioning from a rounded morphology to cells with prominent pseudopodia and protrusions^37–39^. To determine whether mechanical confinement alone is sufficient to induce a comparable phenotypic transition, RAW264.7 cells were confined at heights of 20 μm, 6 μm, and 3 μm (Fig. 1g, supplementary video 1). We found that cells largely retained a rounded morphology comparable to controls under 20 μm slight confinement or 6 μm moderate confinement (Fig. 1g, supplementary video 1). In striking contrast, 3 μm strong confinement triggered a progressive transformation, with cells adopting spindle-shaped bodies and forming dynamic protrusions within 12–24 hours (Fig. 1g and Extended Data Fig. 1d, supplementary video 1). Quantitative analysis showed that ∼30% of cells underwent this transformation after 24 hours of strong confinement (Fig. 1h,i). In parallel, strong confinement markedly enhanced cell motility, which peaked at 12 hours post-confinement (Extended Data Fig. 1e-h). Similar confinement-induced morphological changes were also observed in the human monocytic cell line THP-1, resembling those induced during phorbol 12-myristate 13-acetate (PMA) – induced macrophage differentiation, a process driven by protein kinase C (PKC) activation (Extended Data Fig. 1i–k). Strikingly, transformed cells displayed robust phagocytic activity, efficiently engulfing nearby apoptotic debris (Fig. 1j, supplementary video 2), a functional hallmark typically associated with mature macrophages. Collectively, these observations prompted the hypothesis that long-term strong mechanical confinement may be sufficient to drive monocyte-lineage cells toward macrophage differentiation (Fig. 1k).

### 2.2 Long-term confinement drives RAW264.7 cells toward an activated macrophage-like phenotype

To elucidate the molecular basis of confinement-induced macrophage differentiation, we performed RNA sequencing (RNA-seq) on RAW264.7 cells after 24 hours of strong confinement. This analysis identified 2,550 upregulated and 3,173 downregulated genes (Fig. 2a). Upregulated transcripts included mechanosensitive channels (*Tmem63a/b*) and key macrophage regulators (*Cebpb, Rbpj, Ccl5*). Gene Ontology (GO) enrichment analysis indicated that confinement promoted macrophage-associated processes such as amoeboid migration, antigen presentation, and phagocytosis, while downregulated processes included DNA replication and damage response (Fig. 2b). Transcript-per-million (TPM) analysis confirmed elevated expression of macrophage maturation markers, including *Adgre1*, *Itgam*, and *Cd86* (Fig. 2c and Extended Data Fig. 2a). These findings were further validated by qPCR, which showed increased transcription of macrophage-associated cytokines—IL-1β, TNF-α, and iNos—after 24 hours of strong confinement (Extended Data Fig. 2b).

**Figure 2.**
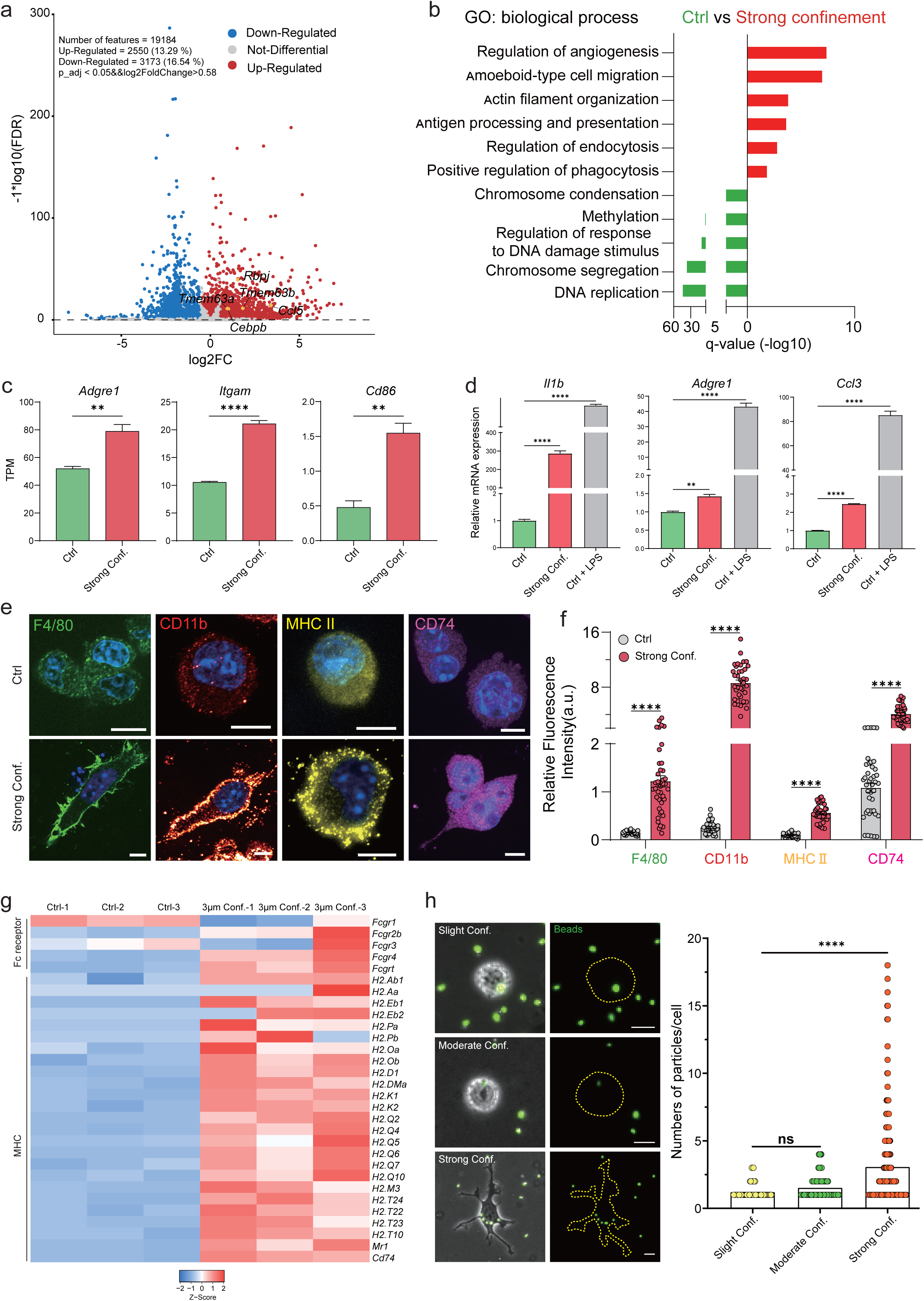
Strong confinement reprograms transcriptional and functional states in RAW264.7 cells. a. Volcano plot showing differentially expressed genes between strong confinement and control groups. b. Gene Ontology enrichment analysis of differentially expressed genes in the strong confinement versus control comparison. c. Quantification of representative transcript levels (TPM) of Adgre1, Itgam, and Cd86 from the RNA-seq dataset. Data are shown as mean ± s.e.m. (*N* = 3 independent experiments). Statistical significance was determined by a two-tailed unpaired Student’s *t*-test; \**P* < 0.01, \*\*\**P* < 0.0001. d. qPCR validation of Il1b, Adgre1, and Ccl3 expression in the control, strong confinement, and LPS-treated groups. Data are shown as mean ± s.e.m. (*N* = 3 independent experiments). Statistical significance was determined by a two-tailed unpaired Student’s *t*-test; \**P* < 0.01, \*\**P* < 0.001, \*\*\**P* < 0.0001. e.f Immunofluorescence staining (e) and fluorescence intensity quantification (f) of macrophage markers in the strong confinement and untreated control groups. F4/80 (green), CD11b (red), MHC II (yellow), and CD74 (magenta) illustrate confinement-induced changes in macrophage identity and activation. Data are shown as mean ± s.e.m. (*N* = 3 independent experiments, F4/80: n = 36 cells in Ctrl and n = 42 cells in Strong Conf.; CD11b: n = 41 cells in Ctrl and n = 38 cells in Strong Conf.; MHC II: n = 47 cells in Ctrl and n = 39 cells in Strong Conf.; CD74: n = 42 cells in Ctrl and n = 40 cells in Strong Conf.). Data were analyzed by a two-tailed unpaired Student’s t-test; ****P < 0.0001. Scale bar, 10 μm. g. Heat map showing expression levels of Fc receptor–related genes and MHC-associated genes in the strong confinement (3 μm) versus untreated control groups. h. Representative images of fluorescent bead phagocytosis and quantification of phagocytic efficiency under Slight Conf. (20 μm), Moderate Conf. (6 μm), and Strong Conf. (3 μm). Data are shown as mean ± s.e.m. (*N* = 3 independent experiments; *n* = 84 cells for Slight Conf., *n* = 79 cells for Moderate Conf., and *n* = 207 cells for Strong Conf.). Statistical significance was determined by a two-tailed unpaired Student’s *t*-test; ns, not significant; \*\*\**P* < 0.0001. Scale bar, 10 μm.

For benchmarking, mechanically confined cells were compared with LPS-activated macrophages. Both conditions elicited robust upregulation of *Adgre1*, *Itgam*, *Il1b*, *Nos2*, *Tnf*, and *Ccl3* (Fig. 2d and Extended Data Fig. 2c), indicating that mechanical stress can recapitulate aspects of classical macrophage activation. Immunofluorescence further confirmed elevated expression of macrophage marker F4/80, CD11b, MHCII, and CD74 in confined RAW264.7 cells (Fig. 2e,f). In addition, gene sets related to Fc receptor signaling and antigen processing and presentation were significantly upregulated following long-term strong confinement (Fig. 2g). Functionally, 3 μm confined cells exhibited markedly enhanced phagocytic capacity, as demonstrated by increased uptake of fluorescent beads (Fig. 2h). Collectively, these findings demonstrate that long-term mechanical confinement initiates a transcriptional and functional program characteristic of macrophage differentiation.

### 2.3 Mechanical confinement regulates primary bone marrow–derived monocytes into macrophages

Having established that mechanical confinement drives macrophage-like differentiation in monocyte-lineage cells, we next asked whether this response is conserved in primary monocytes to determine its physiological relevance. We applied strong mechanical confinement to primary murine bone marrow–derived monocytes (mMonocytes) and human umbilical cord blood–derived monocytes (hMonocytes) (Fig. 3a). Both cell types adopted elongated, spindle-like morphologies with prominent protrusions following prolonged confinement (Fig. 3b,c), recapitulating the responses observed in RAW264.7 and THP-1 cells. These results demonstrate that confinement-induced macrophage differentiation is not restricted to transformed cell lines but represents a conserved, intrinsic property of monocytes.

**Figure 3.**
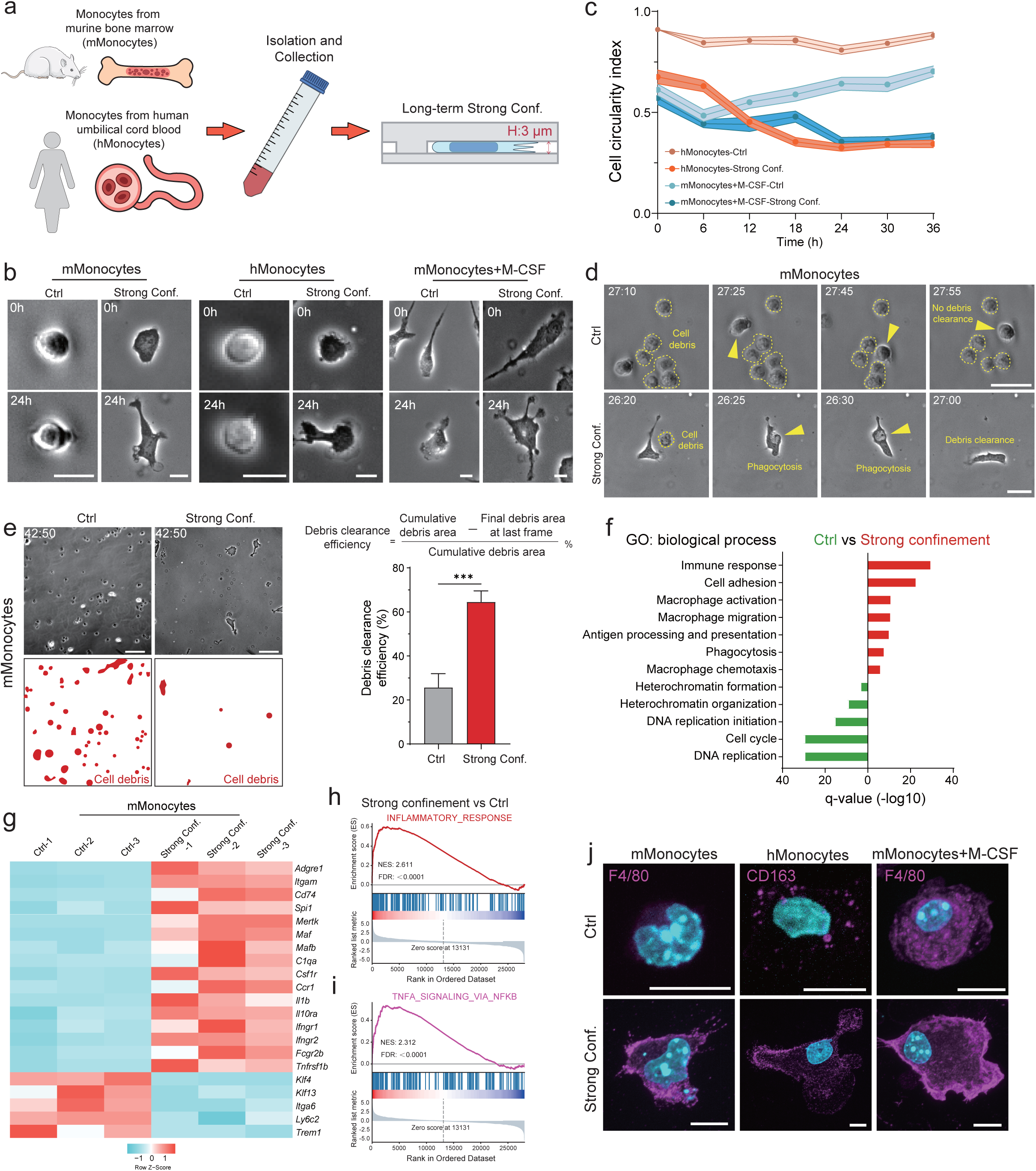
Long-term strong confinement regulates monocyte-to-macrophage differentiation in primary monocytes. a. Schematic illustration of strong confinement applied to freshly isolated mouse bone marrow–derived monocytes (mMonocytes) and human umbilical cord blood–derived monocytes (hMonocytes). b,c. Representative images (b) and cell circularity index (c) of mMonocytes, hMonocytes, and M-CSF – treated mMonocytes (20 ng/mL) cultured under untreated control (no confinement) or 3 μm strong confinement for long-term incubation. Scale bar, 10 μm. d. Representative images showing strong confinement–activated phagocytosis of cellular debris in mMonocytes. Scale bar, 20 μm. e. Representative images (left) and debris clearance efficiency (right) of mMonocytes under untreated control and strong confinement during ∼48 h live tracking. Data are shown as mean ± s.e.m. (*N* > 3 independent experiments). Statistical significance was determined by a two-tailed unpaired Student’s *t*-test; \*\*\**P* < 0.001. Scale bar, 50 μm. f. GO enrichment analysis of differentially expressed genes in strong confinement versus untreated control mMonocytes. g. Heat map showing the expression levels of macrophage activation – associated genes in mMonocytes under strong confinement versus untreated control conditions. h,i. GSEA of INFLAMMATORY_RESPONSE (h) and TNFA_SIGNALING_VIA_NFKB (i) gene sets in strong confinement versus untreated control mMonocytes. j. Representative images of macrophage markers F4/80 and CD163 in mMonocytes, hMonocytes, and M-CSF–treated mMonocytes under control and strong confinement conditions. Scale bar, 10 μm.

In the canonical biochemical pathway, macrophage colony-stimulating factor (M-CSF) drives monocyte-to-macrophage differentiation over 5–7 days^40,41^. To test whether mechanical confinement cooperates with biochemical cues, we treated mMonocytes with low-dose M-CSF (20 ng/mL) for 2 days followed by strong confinement for 1 day. Combined treatment accelerated differentiation, transforming mMonocytes from a spindle-like to a protrusive macrophage-like phenotype compared with non-confined cells (Fig. 3b,c, supplementary video 3), indicating that mechanical and biochemical signals synergistically regulate macrophage maturation.

Monocytes typically possess crescent-shaped nuclei (Extended Data Fig. 3a,b, the region outlined in red). Following long-term confinement, both mMonocytes and hMonocytes displayed oval or spindle-shaped nuclei, consistent with nuclear deformation in response to mechanical stress (Extended Data Fig. 3a,b, the region outlined in blue). Confinement also enhanced phagocytic capacity: unlike unconfined controls, confined cells efficiently engulfed surrounding apoptotic debris (Fig. 3d and Extended Data Fig. 3c, supplementary video 4,5). In contrast, lineage-negative (Lin⁻) bone marrow cells exhibited morphological changes but minimal phagocytic activity under identical conditions (Extended Data Fig. 3d), underscoring the monocyte-specific nature of mechanically induced phagocytosis. Given the short (∼16–24 h) half-life of mMonocytes, time-lapse imaging initially revealed substantial cell death and debris accumulation under confinement (Fig. 3e, left, supplementary video 6). Strikingly, after ∼42 h, debris was markedly reduced, and quantitative analysis confirmed that prolonged confinement potently activates phagocytosis to drive debris clearance (Fig. 3e, right, supplementary video 6). Consistently, fluorescent bead uptake assays demonstrated robust phagocytic activity under confinement (Extended Data Fig. 3e,f).

To further determine whether confinement activates macrophage-associated gene programs, we compared the transcriptomes of mMonocytes before and after mechanical confinement. Strong confinement broadly upregulated immune-related processes (Fig. 3f). GO analysis revealed enrichment of pathways linked to immune activation and macrophage differentiation, whereas downregulated processes were associated with cell proliferation and heterochromatin assembly—consistent with observations in RAW264.7 cells (Fig. 2b, 3f). At the gene level, confinement upregulated macrophage maturation markers (*Adgre1, Itgam, Mertk, Il1b*) and downregulated monocyte-associated transcripts (*Klf4, Klf13, Itga6, Ly6c2*) (Fig. 3g). Gene Set Enrichment Analysis (GSEA) further confirmed significant enrichment of inflammatory response and TNF signaling pathways (Fig. 3h,i). Immunofluorescence analysis corroborated these findings, showing increased surface expression of macrophage markers F4/80 and CD163 in both murine and human monocytes after confinement, an effect further potentiated by low-dose M-CSF (Fig. 3j and Extended Data Fig. 3g). Moreover, long-term confinement markedly enhanced the formation of actin-rich podosome structures—a hallmark of macrophage cytoskeletal remodeling—in mBMDMs, hUCBDMs, and low-dose M-CSF–treated mBMDMs (Extended Data Fig. 3h,i). Collectively, these data support a role for mechanical confinement in driving monocyte-to-macrophage differentiation and synergizing with biochemical cues to potentiate macrophage maturation and function.

### 2.4 Confinement promotes the differentiation of tissue-resident monocytes into macrophages

To determine whether confinement-driven monocyte-to-macrophage differentiation also occurs in tissue-resident monocytes, we isolated mouse hepatic-resident monocytes (HRMs) (Fig. 4a and Extended Data Fig. 4a,b). Under non-confined conditions, HRMs remained largely rounded with minimal spreading (Fig. 4b,c and Extended Data Fig. 4c, supplementary video 7). In contrast, strong confinement induced pronounced morphological remodeling: cells transitioned into protrusion-rich, octopus-like forms with increased spreading area and reduced circularity, closely mirroring the responses of RAW264.7, mMonocytes, and hMonocytes (Fig. 4b,c and Extended Data Fig. 4c, supplementary video 7). Long-term confinement further promoted the acquisition of macrophage-like phagocytic functions (Fig. 4d, supplementary video 8). Confined mHRMs displayed active migration and dynamic protrusions and rapidly engulfed surrounding cellular debris — behavior absent in unconfined controls (Fig. 4d and Extended Data Fig. 4d, supplementary video 8). This functional activation was accompanied by elevated expression of macrophage markers F4/80 and MHCII (Fig. 4e,f). Notably, the morphology of confined mHRMs closely resembled that of in vivo liver capsular macrophages (LCMs) (Fig. 4g), suggesting that tissue-level mechanical forces may promote monocyte-to-macrophage differentiation in situ.

**Figure 4.**
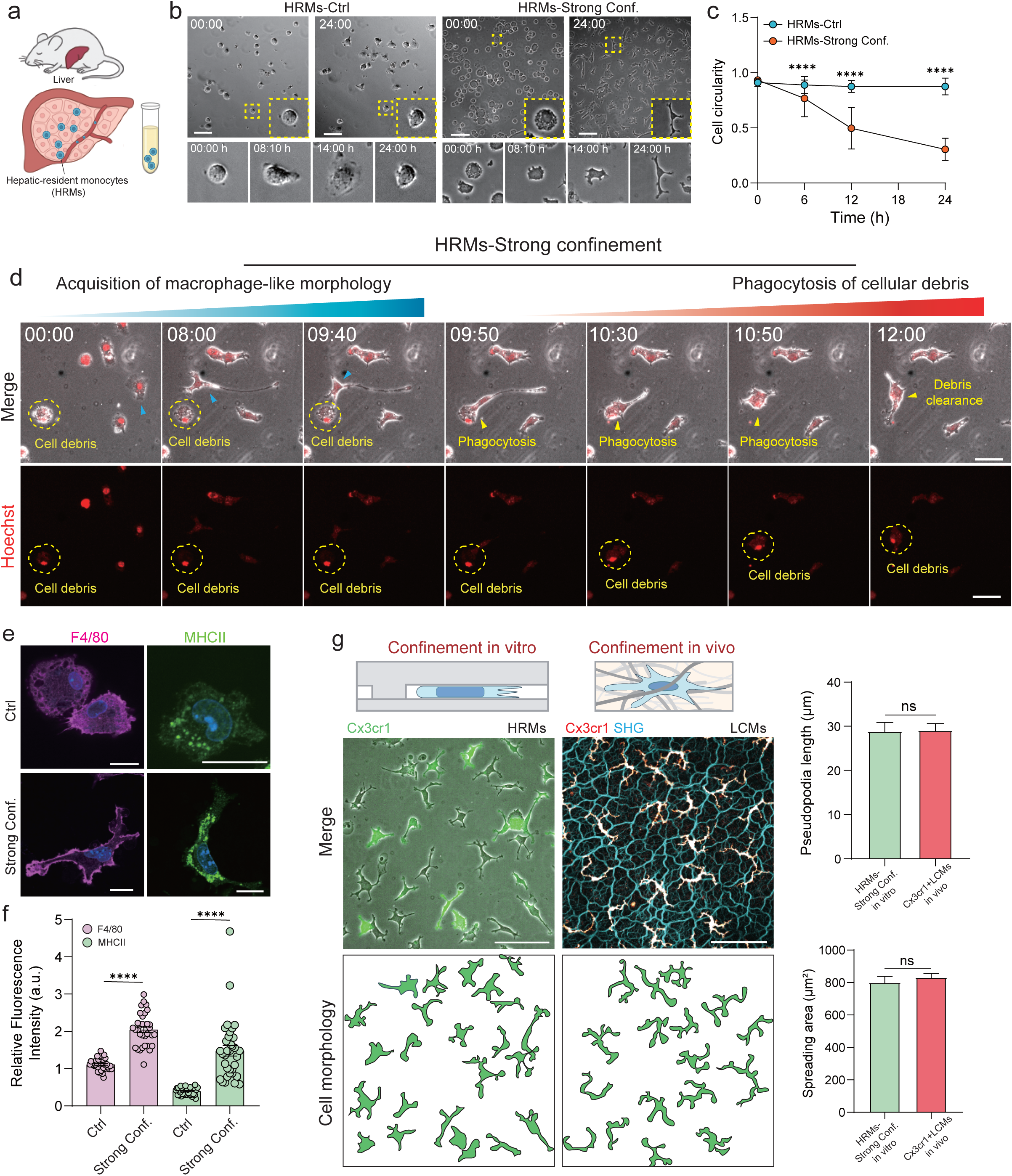
Long-term strong confinement induces macrophage transition in hepatic-resident monocytes. a. Schematic illustration of freshly isolated hepatic-resident monocytes (HRMs) from mouse liver. b,c. Representative images (b) and cell circularity index (c) of HRMs cultured under control (no confinement) or strong confinement (3 μm) during long-term incubation. Data are shown as mean ± s.e.m. (*N* = 3 independent experiments; *n* = 56 cells for HRMs-0 h, *n* = 51 cells for HRMs-6 h, *n* = 51 cells for HRMs-12 h, *n* = 53 cells for HRMs-24 h; *n* = 54 cells for HRMs-strong confinement-0 h, *n* = 54 cells for HRMs-strong confinement-6 h, *n* = 50 cells for HRMs-strong confinement-12 h, and *n* = 52 cells for HRMs-strong confinement-24 h). Statistical significance was determined by a two-tailed unpaired Student’s *t*-test; ****P < 0.0001. Scale bar (HRMs-Ctrl), 50 μm, Scale bar (HRMs-Strong Conf.), 100 μm. d. Representative images showing strong confinement–activated phagocytosis of cellular debris in HRMs. Scale bar, 50 μm. e,f. Representative images (e) and quantitative analysis (f) of macrophage markers F4/80 and MHC II in HRMs under control and strong confinement conditions. (*N* = 3 independent experiments; F4/80: *n* = 36 cells for Ctrl and *n* = 33 cells for strong confinement; MHC II: *n* = 35 cells for Ctrl and *n* = 34 cells for strong confinement). Statistical significance was determined by a two-tailed unpaired Student’s *t*-test; ****P < 0.0001. Scale bar, 20 μm. g. Representative images (left) and quantitative analysis (right) comparing in vitro confinement–induced morphological changes with in vivo–confined *Cx3cr1*⁺ cells. Data are shown as mean ± s.e.m. (*N* = 3 independent experiments; pseudopodia length: *n* = 43 cells for HRMs under strong confinement *in vitro* and *n* = 45 cells for *Cx3cr1*⁺ LCMs *in vivo*; spreading area: *n* = 42 cells for HRMs under strong confinement *in vitro* and *n* = 41 cells for *Cx3cr1*⁺ LCMs *in vivo*). Statistical significance was determined by a two-tailed unpaired Student’s *t*-test; ns, not significant (*P* > 0.05). Scale bar, 100 μm.

Interestingly, releasing cells from confinement after 48 hours revealed partial reversibility: HRMs maintained macrophage-like morphologies for ∼12 hours before gradually reverting to a rounded state (Extended Data Fig. 4e). Together, these findings support a model in which sustained mechanical confinement within tissue microenvironments reprograms monocytes toward macrophage-like states, thereby contributing to local immune remodeling and functional adaptation.

### 2.5 Confinement regulates KDM6B-dependent epigenetic remodeling and macrophage gene activation

The nucleus serves as a central hub for mechanosensation, capable of perceiving mechanical cues and orchestrating downstream adaptive responses^2,42–45^. Under 3 µm confinement, cells exhibited pronounced nuclear deformation, reflecting the strong compression imposed on the nucleus (Fig. 5a and Extended Data Fig. 5a). Quantitative analysis revealed that, relative to controls, confinement markedly flattened nuclei, significantly increasing their maximal projection area (Fig. 5a,b). RNA-seq profiling of mMonocytes and RAW264.7 cells showed that long-term confinement downregulated genes involved in chromatin and heterochromatin assembly (Fig. 2b,5c). These results suggest a hypothesis in which sustained nuclear deformation drives chromatin reorganization and heterochromatin remodeling, thereby reprogramming the transcriptional landscape to fine-tune cellular mechanoadaptation.

**Figure 5.**
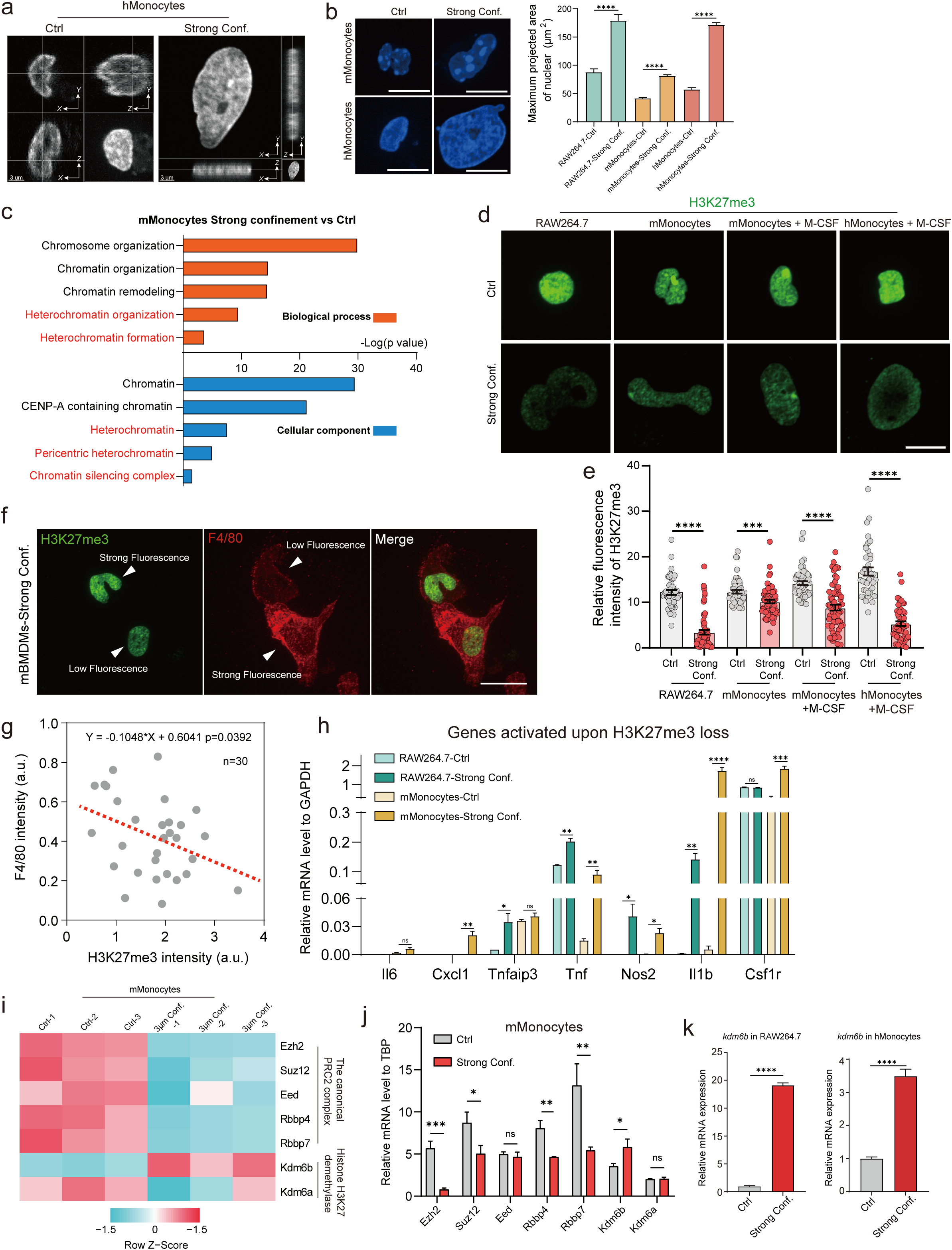
Long-term strong confinement induces heterochromatin remodeling in monocytes. a. Representative images of nuclear morphology in hMonocytes under control and strong confinement conditions. b. Representative images of maximum nuclear projected area in mMonocytes and hMonocytes (left), and quantitative analysis (right) in RAW264.7 cells, mMonocytes, and hMonocytes under control and strong confinement conditions. Data are shown as mean ± s.e.m. (*N* = 3 independent experiments; *n* = 44 cells for RAW264.7-Ctrl, *n* = 61 cells for RAW264.7-Strong Conf.; *n* = 54 cells for mMonocytes-Ctrl, *n* = 68 cells for mMonocytes- Strong Conf.; *n* = 40 cells for hMonocytes-Ctrl, and *n* = 53 cells for hMonocytes- Strong Conf.). Statistical significance was determined by a two-tailed unpaired Student’s *t*-test; ****P < 0.0001. Scale bar, 10 μm. c. GO enrichment analysis of differentially expressed genes associated with chromatin states in strong confinement versus control mMonocytes. d,e. Representative images (d) and relative fluorescence intensity analysis (e) of H3K27me3 in RAW264.7 cells, mMonocytes, M-CSF–treated mMonocytes, and M-CSF–treated hMonocytes under control and strong confinement conditions. Data are shown as mean ± s.e.m. (*N* = 3 independent experiments; RAW264.7: *n* = 44 cells for Ctrl and *n* = 61 cells for Strong Conf.; mMonocytes: *n* = 54 cells for Ctrl and *n* = 68 cells for Strong Conf.; mMonocytes + M-CSF: *n* = 56 cells for Ctrl and *n* = 66 cells for Strong Conf.; hMonocytes + M-CSF: *n* = 40 cells for Ctrl and *n* = 53 cells for Strong Conf.). Statistical significance was determined by a two-tailed unpaired Student’s *t*-test; ***P < 0.001, ****P < 0.0001. Scale bar, 10 μm. f,g. Representative images (f) and correlation analysis (g) of H3K27me3 intensity and the macrophage marker F4/80 in mMonocytes after long-term strong confinement. Scale bar, 20 μm. h. Relative mRNA levels of Il6, Cxcl1, Tnfaip3, Tnf, Nos2, Il1b, and Csf1r in RAW264.7 cells and mMonocytes under control and strong confinement, showing activation of macrophage- and inflammation-associated genes. Data are shown as mean ± s.e.m. (*N* = 3 independent experiments). Statistical significance was determined by a two-tailed unpaired Student’s *t*-test; ns, not significant; *P < 0.05, **P < 0.01, ***P < 0.001, ****P < 0.0001. i,j. Heat map (i) and relative mRNA levels (j) showing expression of Ezh2, Suz12, Eed, Rbbp4, Rbbp7, Kdm6a, and Kdm6b in mMonocytes under control and strong confinement, highlighting confinement-induced alterations in Polycomb complex components and demethylases. Data are shown as mean ± s.e.m. (*N* = 3 independent experiments). Statistical significance was determined by a two-tailed unpaired Student’s *t*-test; ns, not significant; *P < 0.05, **P < 0.01, ***P < 0.001. k. Relative mRNA expression of Kdm6b in RAW264.7 cells and hMonocytes under control and strong confinement, confirming consistent upregulation across species and cell types. Data are shown as mean ± s.e.m. (*N* = 3 independent experiments). Statistical significance was determined by a two-tailed unpaired Student’s *t*-test; ****P < 0.0001.

Among the key regulators of such remodeling are the repressive histone marks H3K9me3 and H3K27me3^46–48^. H3K9me3 is typically associated with constitutive heterochromatin and stable gene silencing, whereas H3K27me3, deposited by the Polycomb repressive complex 2 (PRC2), mediates facultative heterochromatin formation and dynamically represses lineage-specific genes. Together, these modifications safeguard transcriptionally silent chromatin domains and play pivotal roles in fate determination^49–52^. We next examined whether confinement alters these repressive marks. Immunofluorescence revealed that long-term strong confinement induced a pronounced loss of H3K27me3 in mMonocytes (Extended Data Fig. 5b). Consistently, multiple cell types, RAW264.7 cells, mMonocytes, and M-CSF–treated mMonocytes and hMonocytes, showed a robust reduction of H3K27me3 following mechanical stimulation (Fig. 5d,e), whereas H3K9me3 levels remained largely unchanged (Extended Data Fig. 5c,d). Notably, only strong confinement—rather than slight confinement—was sufficient to trigger H3K27me3 erasure (Extended Data Fig. 5c). To monitor H3K27me3 dynamics, we first tracked its reader protein CBX2. Under strong confinement, CBX2 fluorescence progressively decreased, indicating a loss of H3K27me3, whereas CBX2 levels remained stable in unconfined controls (Extended Data Fig. 5e–h). In contrast, the H3K9me3 reader CBX1 showed no noticeable change under strong confinement, serving as an internal control (Extended Data Fig. 5e–h).

Correlation analysis further revealed an inverse relationship between H3K27me3 and macrophage marker F4/80 expression—confined cells with lower H3K27me3 levels exhibited stronger F4/80 expression (Fig. 5f,g), suggesting that confinement-induced H3K27me3 loss may promote macrophage differentiation. Consistent with this, strong confinement also increased H3K27ac, an active enhancer mark that reflects the opening of regulatory chromatin elements and the transcriptional activation of macrophage-associated genes (Extended Data Fig. 5i,j). Confinement-induced erasure of H3K27me3 consequently activated numerous macrophage-associated genes, including *Cxcl1*, *Tnfaip3*, *Tnf*, *Nos2*, *Il1b*, and *Csf1r* (Fig. 5h). Together, these findings identify H3K27me3—but not H3K9me3—as a mechanoresponsive epigenetic mark erased in response to nuclear deformation, linking mechanical confinement to chromatin reprogramming and macrophage differentiation.

Mechanistically, confinement suppressed PRC2 expression while upregulating the H3K27me3 demethylase KDM6B (Fig. 5i-k and Extended Data Fig. 5k,l). This reciprocal regulation drove H3K27me3 erasure, establishing a direct connection between confinement-induced nuclear stress and the epigenetic reprogramming underlying monocyte-to-macrophage differentiation. To validate this pathway, we inhibited KDM6B with GSK-J4 (a cell-permeable inhibitor of the H3K27me3 demethylases KDM6A/B). Under confinement, mechanical stimulation normally activated KDM6B to erase H3K27me3 and induce macrophage differentiation, whereas GSK-J4 treatment restored H3K27me3 levels and suppressed confinement-driven programming (Fig. 6a-d and Extended Data Fig. 6a–c). RNA-seq confirmed that blocking H3K27me3 erasure attenuated expression of macrophage-related genes and downregulated pathways including type II interferon response, antigen presentation, macrophage differentiation, and cytokine production (Fig. 6a,b). Conversely, developmental programs associated with high-H3K27me3 states were upregulated (Fig. 6c). Key macrophage activation genes such as *Adgre1, Itgam, Cx3cr1, Cd74, and Il1b* were markedly downregulated by GSK-J4 (Fig. 6d). Morphologically, GSK-J4 treatment suppressed confinement-induced protrusion formation (Fig. 6e,f), reduced *Itgam* and *Adgre1* expression (Fig. 6g,h), diminished surface F4/80, and impaired cell migration (Fig. 6i and Extended Data Fig. 6d). These results establish KDM6B-mediated H3K27me3 erasure as a critical mechanoresponsive mechanism regulating monocyte-to-macrophage differentiation.

**Figure 6.**
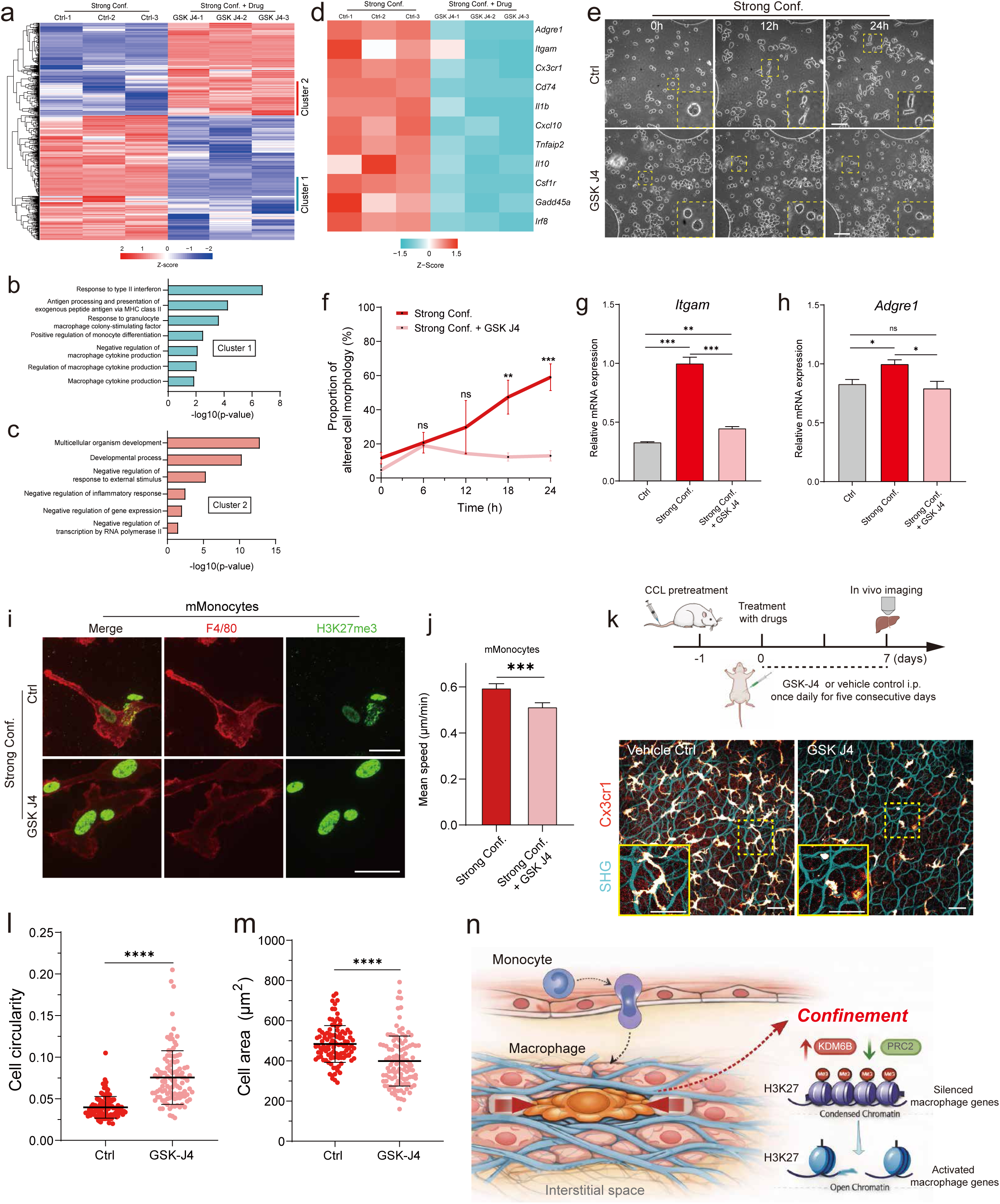
Confinement-activated Kdm6b regulates monocyte-macrophage differentiation. a. Heat map showing gene expression profiles of mMonocytes pretreated with 20 ng/mL M-CSF for 48 h, cultured under strong confinement and treated with DMSO or 10 μM GSKJ4. b,c. GO enrichment analysis of genes from cluster 1 (b) and cluster 2 (c) identified in (a). d. Heat map showing the expression levels of macrophage activation – associated genes in mMonocytes pretreated with 20 ng/mL M-CSF for 48 h, cultured under strong confinement and treated with DMSO or 10 μM GSKJ4. e,f. Representative images (e) and quantitative analysis (f) of cell morphology in mMonocytes under DMSO or GSKJ4-treated strong confinement conditions. Data are shown as mean ± s.e.m. (*N* = 3 independent experiments) Statistical significance was determined by a two-tailed unpaired Student’s *t*-test; ns, not significant; *P < 0.05, **P < 0.01, ***P < 0.001. Scale bar, 100 μm. g,h. qPCR analysis of Itgam (g) and Adgre1 (h) expression in control, strong confinement, and GSKJ4-treated strong confinement groups. Data are shown as mean ± s.e.m. (*N* = 3 independent experiments). Statistical significance was determined by a two-tailed unpaired Student’s *t*-test; ns, not significant; *P < 0.05, **P < 0.01, ***P < 0.001. i,j. Representative fluorescent images (i) showing F4/80 (red) and H3K27me3 (green), and mean migration speed analysis (j) of mMonocytes pretreated with 20 ng/mL M-CSF for 48 h under DMSO or GSKJ4-treated strong confinement conditions. Data are shown as mean ± s.e.m. (*N* = 3 independent experiments; *n* = 248 cells for strong confinement and *n* = 241 cells for strong confinement + GSKJ4). Statistical significance was determined by a two-tailed unpaired Student’s *t*-test; ***P < 0.001. Scale bar, 20 μm. k. Schematic illustration of the in vivo experimental workflow used to assess the effect of GSKJ4 on the maturation of liver capsular macrophages (top), together with representative two-photon intravital fluorescence images (bottom) showing *Cx3cr1* (red–hot) and collagen fibers visualized by second harmonic generation (SHG) signals (cyan). Scale bar, 50 μm. l,m. Quantification of cell circularity (l) and cell area (m) of *Cx3cr1*⁺ liver capsular macrophages (LCMs) *in vivo* under vehicle control or GSKJ4 treatment. Data are shown as mean ± s.e.m. (*N* = 3 independent experiments; *n* = 106 cells for vehicle control and *n* = 110 cells for GSKJ4 treatment). Statistical significance was determined by a two-tailed unpaired Student’s *t*-test; ****P < 0.0001. n. Schematic model illustrating confinement-driven monocyte-to-macrophage differentiation mediated by Kdm6b-dependent epigenetic remodeling.

Finally, to test this mechanism in vivo, we depleted liver capsular macrophages (LCMs) with clodronate liposomes, enabling circulating monocytes to repopulate the capsule (Fig. 6k). In untreated mice, infiltrating monocytes gradually acquired octopus-like morphologies indicative of macrophage maturation (Fig. 6k). In contrast, GSK-J4–treated mice showed markedly impaired LCM replenishment, and residual cells remained rounded and immature, as visualized by two-photon imaging in vivo (Fig. 6k-m). These in vivo findings mirror our in vitro observations and demonstrate that KDM6B-dependent H3K27me3 erasure is not only sufficient but also required for confinement-driven macrophage differentiation, establishing mechanical confinement as a physiologically essential regulator of monocyte fate in tissue environments.

Collectively, our findings reveal that strong mechanical confinement represents a potent and previously underappreciated physical cue that regulates monocyte-to-macrophage differentiation (Fig. 6n). We further identify a mechano-epigenetic regulatory axis whereby long-term confinement is associated with activation of KDM6B and the removal of H3K27me3, thereby facilitating chromatin accessibility and transcriptional reprogramming required for macrophage identity acquisition (Fig. 6n). Together, these observations highlight a previously unrecognized dimension of immune regulation governed by tissue mechanics and underscore the potential of mechanical confinement as an engineering-compatible cue for generating tailored macrophage phenotypes, with implications for immune modulation and macrophage-based therapeutic strategies.

## Discussion

Mechanical confinement is a pervasive physical feature of tissue environments, yet its role in immune cell fate regulation has remained poorly defined. Here, we discover that sustained strong confinement can drive monocyte-to-macrophage differentiation, inducing a pronounced cell shape transition from a rounded state to a highly spread, protrusion-rich macrophage-like phenotype. Cell shape remodeling is a sensitive and easily quantifiable readout of cellular phenotype transitions. In vivo, within the confined microenvironment of the liver capsule, mature macrophages exhibited a highly extended, octopus-like morphology. We therefore sought to understand why rounded monocytes undergo such pronounced cell shape remodeling during their differentiation into macrophages within confined tissue microenvironments, and to determine whether mechanical confinement represents a critical driver of this process. Strikingly, a similar morphological remodeling was recapitulated in vitro when monocytes were subjected to strong mechanical confinement at a height of 3 μm. This confinement-induced cell shape remodeling was accompanied by upregulation of macrophage marker expression and an enhanced capacity for cellular debris phagocytosis, supporting a linkage between mechanical confinement, cell shape remodeling, and macrophage-like functional acquisition. Notably, comparable confinement-dependent phenotypic plasticity has also been observed in other cell types. Under similar 3 μm confinement, melanoma cells undergo a neuron-like phenotypic transition that modulates their proliferative and invasive states to adapt to mechanical stresses encountered during metastasis^18^. Likewise, transient confinement has been shown to promote neuronal reprogramming in fibroblasts^15^. Together, these observations identify mechanical confinement as an instructive physical cue capable of shaping cell fate decisions across diverse biological contexts. Furthermore, the combination of M-CSF–mediated biochemical signaling and confinement-induced mechanical cues resulted in augmented macrophage functional activation, highlighting that mechanical and biochemical inputs act synergistically to regulate cell shape remodeling and determine immune cell fate. In contrast, under non-adhesive conditions, as demonstrated in our previous work on tumor cells, 3 μm mechanical confinement induces a rapidly migrating A2 phenotype, thereby enabling efficient traversal through confined microenvironments^11^. In addition, confinement within dynamically sliding hydrogels has been shown to induce cell tumbling behaviors that promote chondrogenic differentiation^29^. Together, these findings underscore that distinct mechanical contexts of confinement dictate divergent mechanoadaptive strategies, cell shape remodeling, and cell fate decisions.

Furthermore, we identified KDM6B as a key epigenetic responder to confinement-induced monocyte-to-macrophage differentiation. Sustained strong confinement suppresses components of the PRC2 complex while concomitantly upregulating KDM6B, resulting in selective removal of H3K27me3 and activation of macrophage-associated transcriptional programs. These findings suggested that the epigenome serves as a dynamic interface linking the physical microenvironment to cell identity. Previous studies have highlighted the nucleus as a rapid mechanical “ruler” that senses spatial confinement through nuclear envelope stretching, triggering calcium release and cPLA2 recruitment within minutes to promote fast migratory responses^42,44,53^. In line with this rapid mechanosensing paradigm, transient confinement has been shown to induce short-term changes in chromatin states, such as alterations in H3K9me3, thereby facilitating fibroblast-to-neuron differentiation^15^. However, how long-term mechanical confinement regulates cell fate decisions remains poorly understood. In this work, we found that prolonged confinement activates KDM6B-mediated H3K27me3 erasure and drives macrophage-associated transcriptional programs that support long-term mechanoadaptation. Whether heterochromatin modification is altered in response to mechanical stimulation appears to depend on both the duration of force exposure and the cellular context. Transient mechanical compression does not markedly change H3K27me3 levels in fibroblasts^15^, whereas confinement within sliding hydrogel systems preserves H3K27me3 in mesenchymal stem cells, thereby maintaining their osteogenic differentiation potential^29^. Recent studies suggest that mechanical confinement is associated with regulation of the DNA-binding protein HMGB2 in melanoma cells, prolonging its contact time with chromatin and coinciding with increased metastatic propensity^18^. In addition, mesenchymal stem cells undergoing long-term confined migration exhibit elevated H3K27ac levels that correlate with enhanced osteogenic differentiation^16^. Collectively, these studies, together with our findings, indicate that confinement-induced chromatin remodeling acts as a fundamental mechanism to regulate cell fate decisions. Despite these advances, key questions remain. How sustained nuclear stress is transduced into KDM6B induction remains unknown. Whether confinement activates specific mechanosensitive transcription factors, alters chromatin accessibility at the Kdm6b locus, or reshapes nucleosome organization are important directions for future investigation. Addressing these questions will be essential for understanding how persistent mechanical cues are encoded into stable epigenetic states that govern immune cell fate.

Beyond establishing mechanical confinement as a potent instructive cue for macrophage reprogramming, our findings suggest new opportunities for mechanobiology-guided immunotherapies. Macrophages are highly abundant in tumors and capable of infiltrating dense, fibrotic, and spatially constrained niches, making them an attractive vehicle for next-generation cancer therapies such as chimeric antigen receptor macrophages (CAR-M)^54,55^. In our study, confinement-programmed macrophages displayed markedly enhanced phagocytic capacity and efficiently cleared surrounding cellular debris — mirroring the functional logic of CAR-M, where anti-CD19/CD22/HER2 CARs encoding CD3ζ or CD3ζ–TIR intracellular domains boost tumor-cell recognition and CAR-driven engulfment^56–58^. Importantly, engineered macrophages naturally maintain a pro-inflammatory phenotype, a feature that strengthens phagocytosis, enhances tumoricidal activity, and promotes antigen presentation to amplify downstream T-cell responses^57,58^. Many hallmark pro-inflammatory and interferon-response genes (*IL1A, IL1B, IL6, IL12A, TNF, IFIT1, ISG15, IFITM1*^57,58^) were likewise robustly upregulated in confinement-programmed macrophages, indicating that strong confinement pushes macrophages toward an immunostimulatory state. Thus, mechanical confinement could serve as a non-genetic activation modality or be combined with CAR constructs to further enhance CAR-M functionality. Together, these insights establish mechanical confinement as a powerful and complementary axis for controlling macrophage identity and function, with broad implications for immunity, tissue biology, and cell-based therapies.

## Methods

### Reagents

Deionized (DI) water (18.2 MΩ· cm) was produced using a Milli-Q system (Millipore, Bedford, MA, USA) and used for all experiments. Fibronectin, GSK-J4 hydrochloride, phorbol 12-myristate 13-acetate (PMA), lipopolysaccharide (LPS; Escherichia coli O111:B4), and mouse M-CSF protein were purchased from MedChemExpress. The following antibodies were obtained from BioLegend: APC anti-mouse F4/80, PE anti-mouse/human CD11b, Alexa Fluor® 647 anti-mouse CD74 (CLIP), APC anti-human CD163, Pacific Blue™ anti-mouse/human CD11b, Brilliant Violet 510™ anti-mouse CD45, FITC anti-mouse Ly-6C, and APC anti-mouse Ly-6G antibodies. Alexa Fluor™ 647 Phalloidin and FITC anti-mouse MHC II (I-A/I-E) antibodies were purchased from Thermo Fisher Scientific (Invitrogen). Amine-modified fluorescent yellow–green polystyrene latex beads and Hoechst 33342 were purchased from Sigma-Aldrich. Six-well glass-bottom plates and 35-mm glass-bottom dishes were obtained from Cellvis. The histone H3K9me3 polyclonal antibody was purchased from Active Motif, and the tri-methyl-histone H3 (Lys27) rabbit monoclonal antibody (C36B11) was obtained from Cell Signaling Technology (CST). The Monocyte Isolation Kit (mouse bone marrow), Pan Monocyte Isolation Kit (human), and Lineage Cell Depletion Kit (mouse) were purchased from Miltenyi Biotec. Clodronate liposomes was purchased from Yeasen.

### Animal

*Cx3cr1*^GFP/+^ mice was gifts from Prof. Florent Ginhoux. All animals were maintained under specific pathogen free (SPF) conditions at the Center for Excellence in Molecular Cell Science, Chinese Academy of Sciences, with a 12-hour light/dark cycle and ad libitum access to standard chow. All procedures were approved by the Institutional Animal Care and Use Committee (IACUC) of the Shanghai Institute of Biochemistry and Cell Biology. Mice aged 8–12 weeks were used for most experiments.

### Cell Culture

RAW264.7 cells (kindly provided by Fei Lan, Fudan University) and RAW264.7 Myrpalm-mEmerald cells were cultured in Dulbecco’s Modified Eagle Medium (DMEM; GIBCO) supplemented with 10% fetal bovine serum (FBS; ExCell Bio) and 100 U/mL penicillin–streptomycin (GIBCO) at 37 °C in a humidified atmosphere containing 5% CO₂.

mMonocytes, mMonocytes treated with 20 ng/mL mouse M-CSF for 2 days, bone marrow-derived Lin⁻ cells, hMonocytes, hMonocytes treated with 20 ng/mL human M-CSF for 2 days, THP-1 cells, and HRMs were maintained in RPMI 1640 medium (GIBCO) supplemented with 10% FBS (ExCell Bio) and 100 U/mL penicillin–streptomycin (GIBCO) under the same culture conditions. Prior to live-cell imaging, all samples were incubated with 200 ng/mL Hoechst 33342 at 37 °C for 1 h to label cell nuclei.

### Isolation of mMonocytes, Bone Marrow-Derived Lin⁻ Cells, hMonocytes

Murine monocytes (mMonocytes) and bone marrow–derived Lin⁻ cells were isolated from the bone marrow of 8-week-old male C57BL/6 mice. Human monocytes (hMonocytes) were obtained from human umbilical cord blood samples. All cell isolation procedures were performed according to the manufacturers’ instructions provided with the respective isolation kits (Miltenyi Biotec; Order Nos. 130-096-537, 130-090-858, and 130-100-629). The procedure for mMonocyte isolation is described below as a representative example; the isolation of hMonocytes and bone marrow–derived Lin⁻ cells was performed using similar procedures, with differences only in the reagents applied.

### Preparation of murine bone marrow cells

Murine bone marrow cells were harvested from the femurs and tibias by flushing the marrow cavity with cold buffer using a syringe fitted with a 26-gauge needle. The resulting cell suspension was gently dissociated by repeated pipetting and passed through a 30 μm nylon mesh to remove cell clumps. Cells were washed by centrifugation at 300 × g for 10 min at 4 °C, and the supernatant was discarded.

### Magnetic labeling for mMonocyte isolation

The cell pellet was resuspended in 175 μL of buffer per 5 × 10⁷ total cells. Fc receptor (FcR) Blocking Reagent (25 μL per 5 × 10⁷ cells) was added, mixed thoroughly, and incubated briefly. Subsequently, 50 μL of Monocyte Biotin–Antibody Cocktail per 5 × 10⁷ cells was added, mixed well, and incubated for 5 min at 4 °C.

Cells were then washed with 10 mL of buffer per 5 × 10⁷ cells and centrifuged at 300 × g for 10 min at 4 °C. After complete removal of the supernatant, the cell pellet was resuspended in 400 μL of buffer per 5 × 10⁷ cells, followed by addition of 100 μL of Anti-Biotin MicroBeads per 5 × 10⁷ cells. The suspension was mixed thoroughly and incubated for 10 min at 4 °C.

### Magnetic separation

LS Columns were placed in the magnetic field of a MACS Separator and equilibrated with 3 mL of buffer. The labeled cell suspension was applied onto the column, and the flow-through containing unlabeled cells was collected as the enriched monocyte fraction. The column was washed three times with 3 mL of buffer, and all flow-through fractions were combined to obtain purified mMonocytes.

### Hepatocyte-conditioned expansion and purification of Hepatic-resident monocytes

Primary hepatocytes were isolated from 2-month-old mice using a two-step collagenase perfusion method as previously described^59^. Briefly, mice were anesthetized with sodium pentobarbital (60 mg/kg), and the liver was perfused via the inferior vena cava using a peristaltic pump (LONGER BT100-2J). Perfusion was initiated with a calcium-free buffer containing 137 mM NaCl, 5.37 mM KCl, 0.5 mM NaH₂PO₄·2H₂O, 0.42 mM Na₂HPO₄·12H₂O, 4.17 mM NaHCO₃, 0.5 mM EGTA, and 5 mM glucose (pH 7.4), followed by enzymatic digestion with a buffer composed of 137 mM NaCl, 5.37 mM KCl, 0.5 mM NaH₂PO₄·2H₂O, 0.42 mM Na₂HPO₄·12H₂O, 4.17 mM NaHCO₃, 10 mM HEPES, 1 mM CaCl₂, and 0.3 mg/mL Collagenase IV (Sigma-Aldrich, #C5138-1G), pre-warmed to 37 °C (pH 7.4). The hepatic portal vein was clipped to prevent backflow and ensure uniform perfusion, and gentle manual compression of the liver was applied to facilitate even enzyme distribution.

Following perfusion, livers were excised and gently dissociated in a Petri dish. The resulting cell suspension was passed through a 150-mesh nylon cell strainer and centrifuged at 50 × g for 3 min to pellet hepatocytes. The supernatant, containing non-parenchymal cells, was collected and centrifuged at 300 × g for 5 min. Hepatocyte pellets were further purified by centrifugation through a 45% Percoll gradient (Sigma-Aldrich, #P1644) at 50 × g for 10 min. Primary co-cultures were initially washed with PBS and incubated with 0.25% trypsin at 37 °C for 5 min. The culture dish was gently agitated to selectively detach hepatocytes, which appeared as white cellular aggregates. Detached hepatocytes were removed by rinsing with 1× PBS. This procedure was repeated as necessary until the majority of hepatocytes were eliminated, leaving predominantly smaller, ovoid phagocytic cells adherent to the culture surface.The remaining adherent cells were subsequently digested with Accutase (BD Biosciences, #561527) at room temperature for 5 min. Enzymatic digestion was quenched by the addition of DMEM supplemented with 10% FBS. Hepatic-resident monocyte colonies were gently dislodged by pipetting and collected by centrifugation at 450 × g for 5 min at 4 °C. The supernatant was discarded, and the cell pellet was resuspended in serum-free DMEM for downstream applications.

### Cell Confiner Fabrication

The cell confiner device was developed as a modified design based on a conventional six-well plate culture system. It consisted of a customized six-well plate lid, large PDMS pillars, and small PDMS pillars. A 100-μm thick Gel-film sheet (Gel-Pak) was affixed to the inner surface of the plate lid, and large PDMS pillars were bonded to the sheet so that each pillar was positioned at the center of a well. When the lid was closed, the pillars pressed confining slides onto the culture substrate, thereby physically restricting the cells.

### Fabrication of large PDMS pillars

Large PDMS pillars were fabricated by pouring a PDMS mixture (base:curing agent = 35:1, Sylgard 184, Dow Corning) into a custom-made mold, followed by degassing under vacuum. The mixture was cured for one month at room temperature and demolded using a small amount of isopropanol to facilitate release. Owing to the low curing agent ratio, the resulting PDMS pillars were soft and adhesive, which ensured firm attachment to the lid after cleaning.

### Fabrication of small PDMS pillars on glass slides (confining slides)

Small PDMS pillars were fabricated by standard photolithography. Briefly, an SU8-2005 photoresist (MicroChem) was spin-coated onto a silicon wafer and patterned to form a regular array of holes (diameter: 440 μm, spacing: 1 mm), following the manufacturer’s protocol. The mold was silanized with trimethylchlorosilane (TMCS) vapor for 3 min. A drop of PDMS mixture (base:curing agent = 8:1) was then poured into the SU8 mold, and a Φ12 mm coverslip, freshly plasma-activated for 2 min (Harrick Plasma, Ithaca, NY, USA), was gently pressed onto the PDMS drop to form a thin residual PDMS layer. After baking at 95 °C for 15 min on a hot plate, excess PDMS was removed. The glass slide with PDMS pillars was detached by adding a drop of isopropanol and gently lifting the slide with a razor blade.

### Assembly and preparation

Prior to use, the modified six-well plate lid with large PDMS pillars was sterilized with 70% ethanol. Confining slides were coated with fibronectin (25 μg/mL in PBS) for 1 h and equilibrated in culture medium. During assembly, the large PDMS pillars on the lid held the confining slides in place. Upon closing the lid, the pillars pressed the confining slides against the culture substrate, creating a confined microenvironment for cell culture.

### Drug Treatment

RAW264.7 cells or mMonocytes were seeded at a density of 5 × 10⁵ cells/mL on six-well glass-bottom plates coated with 25 μg/mL fibronectin. Cells were treated with 10 μM GSK-J4 for 4 h prior to confinement or non-confinement, and the drug was maintained in the culture medium throughout the 24-h confinement and image acquisition period. For in vivo experiments, GSK-J4 was dissolved in 10% DMSO, 40% PEG300, 5% Tween-80, and 45% saline. Six-week-old male *Cx3cr1*^GFP/+^ mice received a single intravenous injection of 200 μL clodronate liposomes to deplete tissue macrophages, followed by intraperitoneal administration of GSK-J4 (1 mg/kg) or vehicle once daily for 5 of 7 days. LCMs were then imaged by two-photon intravital microscopy.

### Live-cell imaging

Time-lapse imaging was performed using 20×, 40×, or 100× objectives on one of the following imaging systems: a DMi8 inverted microscope (Leica) equipped with an ORCA-Flash4.0 V3 digital CMOS camera (Hamamatsu) and controlled by MetaMorph software (Universal Imaging); a Nikon Eclipse Ti2 microscope (Nikon) equipped with an ORCA-Flash4.0 V3 digital CMOS camera (Hamamatsu) and controlled by NIS-Elements software (Nikon); or a spinning-disk confocal microscope (Yokogawa CSU-W1, Nikon) equipped with a Prime 95B sCMOS camera (Teledyne Photometrics) and controlled by NIS-Elements software (Nikon).

All live-cell imaging experiments were conducted in a temperature- and CO₂-controlled cage incubator (Okolab). Image processing and quantitative analyses were performed using FIJI (ImageJ) and Imaris software.

### Two-photon microscopy

Mice were anesthetized with sodium pentobarbital (50 mg/kg) and placed in a supine position on a thermostatically controlled heating pad maintained at 37 °C. Animals were secured with medical adhesive tape. To minimize motion artifacts during imaging, the abdominal skin and musculature were carefully resected, and a gentle negative-pressure stabilization system was applied to immobilize the liver.

The animal, together with the heating setup, was positioned on the stage of an Olympus FVMPE-RS two-photon microscope. A 25× water-immersion objective lens (NA 1.05) was oriented perpendicular to the region of interest. The liver was gently stabilized against a glass coverslip using mild suction, and 1× PBS was used as the immersion medium between the objective and the tissue.

Imaging parameters were configured using FV-SW software. A MAITAI EHP DeepSee-OL laser was tuned to 870 nm to excite both second harmonic generation (SHG) signals and GFP fluorescence. For volumetric imaging, z-stacks were acquired from the liver capsule to a depth of 60–80 μm, with optical sections collected at 2 μm intervals.

### Phagocytosis assay

FITC-labeled microspheres (Sigma-Aldrich, #L1030) were preconditioned by incubating 1 μL of beads in 100 μL of fetal bovine serum (FBS) at 37 °C for 1 h to promote protein coating. The beads were then pelleted by centrifugation at 12,000 rpm for 3 min, washed once, and resuspended in 100 μL of DMEM. The prepared microspheres were subsequently added to the cell culture for phagocytosis assays.

### Immunofluorescence

Cells subjected to 3 μm strong confinement for 24 h were fixed with 4% paraformaldehyde at 4 °C for 2 days, followed by permeabilization with 0.25% Triton X-100 in PBS at 4 °C for 1 day. After washing with PBS, samples were blocked with 3% bovine serum albumin (BSA) in PBS at 4 °C for 1 day. Cells were then incubated with primary antibodies at 4 °C for 2 days, followed by Alexa Fluor 488-, 594-, or 647-conjugated secondary antibody incubation at 4 °C for 1 day, with extensive PBS washing between steps. Finally, F-actin was stained using Alexa Fluor 594- or 647-conjugated phalloidin, and fluorescence images were acquired using a Nikon fluorescence microscope.

### Analysis of cell trajectories

For cell motility analysis, the nucleus images labeled by Hoechst were tracked using TrackMate plugin in FIJI. Briefly, the calibrated-image sequences were detected by Log detector and were tracked by simple LAP tracker. The spots in tracks statistics were further analyzed by self-wrote code in Matlab to obtain the normalized cell trajectories, mean speed, MSD, and diffusion coefficient. The velocity of migratory cells was calculated according to the displacement of the migratory cells between image sequences divided by timelapse. The mean square displacement (MSD) was calculated as the following Equation (1).

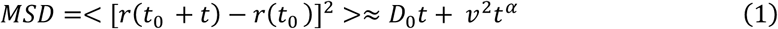

where *v* is the mean speed of migratory cell, *D0* is the thermal diffusion coefficient of the cell, *r* (*t*) is the time dependent position of migratory cell, and *t* is the timelapse of the position.

### RNA Isolation and Library Preparation

RAW264.7 cells or mMonocytes were seeded at a density of 5 × 10⁶ cells/mL on six-well glass-bottom plates coated with 25 μg/mL fibronectin (FN). Cells were subjected to 3 μm strong confinement for 24 h using the cell confiner device. After incubation, the culture medium outside the small PDMS pillars was carefully removed, and both sides of the wells were rinsed with PBS to eliminate non-confined cells. The cell confiner was then gently removed, and the confined cells were collected using TRIzol reagent (Invitrogen).

For RNA extraction, 200 μL of chloroform was added per 1 mL of TRIzol (Thermo Fisher Scientific), followed by vigorous shaking to mix thoroughly. Samples were centrifuged at 12,000 × *g* for 15 min at 4 °C. The aqueous phase was transferred to a new tube, and an equal volume of isopropanol was added. After gentle mixing, the samples were incubated on ice for 10 min and centrifuged again at 12,000 × *g* for 10 min at 4 °C. The supernatant was discarded, and the RNA pellet was washed twice with 75% ethanol, centrifuging at 7,500 × *g* for 5 min each time. After removing the supernatant, the pellet was air-dried at room temperature for 5–10 min and dissolved in RNase-free water. RNA purity and quantification were evaluated using the NanoDrop 2000 spectrophotometer (Thermo Scientific, USA). RNA integrity was assessed using the Agilent 2100 Bioanalyzer (Agilent Technologies, Santa Clara, CA, USA). Then the libraries were constructed using VAHTS Universal V6 RNA-seq Library Prep Kit according to the manufacturer’s instructions. The transcriptome sequencing and analysis were conducted by JMDNA(Shanghai) Bio-Medical Technology Co., Ltd. (Shanghai, China).

### RNA Sequencing and Differentially Expressed Genes Analysis

RNA sequencing libraries were sequenced on a BGI Genomics DNBSEQ-T7 platform, generating 150-bp paired-end reads. Raw FASTQ reads were first processed to remove adaptor sequences and low-quality reads, yielding clean reads for downstream analysis. The clean reads were then aligned to the *Mus musculus* reference genome. Gene expression levels were quantified as FPKM, and read counts for each gene were calculated for subsequent analyses. Principal component analysis (PCA) was performed to assess sample reproducibility and overall variation among groups. Differential expression analysis was conducted to identify significantly changed genes between groups. Genes with an adjusted P value < 0.05 and fold change > 2 or < 0.5 were defined as differentially expressed genes (DEGs). Hierarchical clustering analysis of DEGs was performed to visualize gene expression patterns across samples. A radar plot of the top 30 upregulated or downregulated genes was generated to illustrate representative transcriptional changes. Functional enrichment analyses, including Gene Ontology (GO), KEGG pathway, Reactome, and WikiPathways analyses, were performed for DEGs to identify significantly enriched biological terms and pathways. The enrichment results were visualized using bar plots, chord diagrams, and bubble plots. Gene Set Enrichment Analysis (GSEA) was performed using GSEA software with predefined gene sets. Genes were ranked according to the degree of differential expression between groups, and the enrichment of predefined gene sets at the top or bottom of the ranked list was evaluated.

### Quantitative RT-PCR analysis

Total RNA was extracted using TRIzol reagent (Thermo Fisher Scientific) and quantified by NanoDrop spectrophotometry to assess RNA concentration and purity. Reverse transcription was performed using the MightyScript Plus cDNA Master Mix (Sangon Biotech) according to the manufacturer’s instructions. Quantitative real-time PCR (RT–qPCR) was carried out using a SYBR Green–based PCR master mix on a real-time PCR detection system. Each reaction was performed in a final volume of 20 μL, containing diluted cDNA template, gene-specific primers, and SYBR Green PCR master mix. All samples were analyzed in technical triplicates. The PCR cycling conditions were as follows: an initial denaturation at 95 °C for 3 min, followed by 40 cycles of denaturation at 95 °C for 5 s and annealing/extension at 60 °C for 20 s. Melting curve analysis was performed after amplification to verify product specificity using the following program: 95 °C for 15 s, 60 °C for 1 min, 95 °C for 15 s, and 60 °C for 15 s. Gene expression levels were quantified using primers specific for each target gene, with murine GAPDH or human ACTB (β-actin) used as species-matched internal controls. Ct values were normalized to the corresponding housekeeping gene, and relative expression levels were calculated accordingly. PCR experiments were performed in triplicate and standard deviations calculated and displayed as error bars.

The following primer sequences were used:

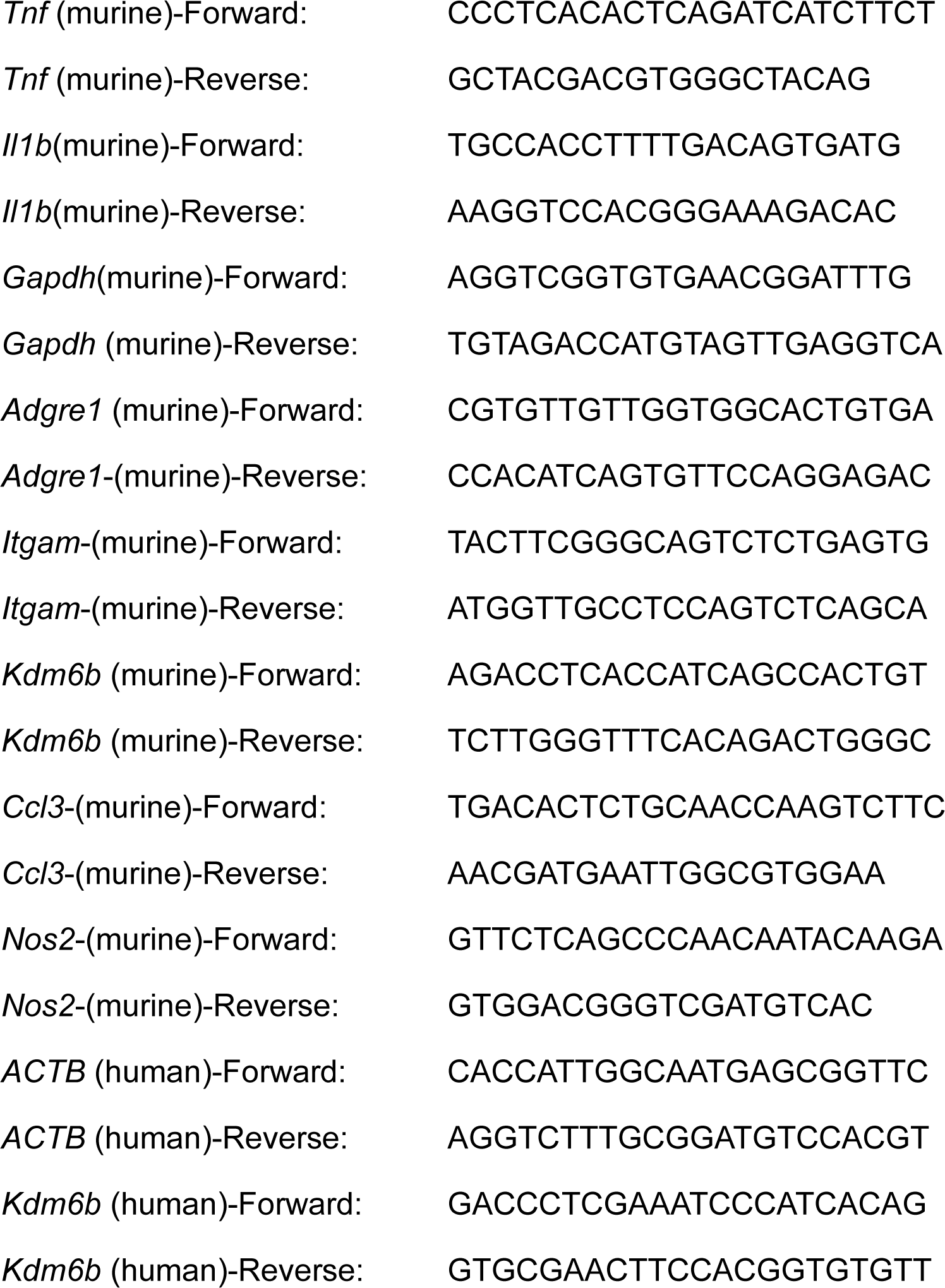

### Statistics

Statistical analyses and data visualization were performed using GraphPad Prism. All data are presented as mean ± s.e.m. The number of biological replicates (*N*) for each experiment is indicated in the corresponding figure legends. Statistical significance is denoted as follows: ns, not significant; *p<0.05, **p<0.01, ***p<0.001, ****p<0.0001 (unpaired t-test).

## Author Contributions

Y.-J.L. conceived the project, supervised the study, and wrote the manuscript. Z.C. and Y.-J.L. supervised the project. W.L., X.-Z.C., and H.Z. performed most of the experiments, analyzed the data, prepared the figures, and wrote the manuscript. X.H. and X.B. provided mMonocytes and hMonocytes. C.F.X., Y.-X.J., and R.-Y.M. collected and analyzed the sequencing data. Y.-T.D., Y.-J.W., M.S., and H.G. assisted with live-cell imaging, animal experiments, and library preparation. G.J. provided methods for cell migration analysis. All authors discussed the results, commented on the manuscript, and approved the final version.

## Acknowledgement

We thank Prof. Fei Lan and Dr. Bowen Rong (Fudan University) for kindly providing RAW264.7 cells and the CBX1-mRuby2 and CBX2-mRuby2 plasmids. We thank Dr. Xiulan Li for her assistance with imaging, as well as the Core Facility of Shanghai Medical College, Fudan University, for technical support. This work was supported by the National Natural Science Foundation of China (Grant No. 22274026 and 32401200), the China Postdoctoral Science Foundation (Grant No. 2024M760695), and the Postdoctoral Fellowship Program of CPSF (Grant No. GZC20240297).

## Extended Data Figure Legends

Extended Data Figure 1. Confinement-induced morphological remodeling in monocyte-lineage cell lines.

a–c. Quantification of cell area (a), solidity (b), and Feret’s diameter (c) of *Cx3cr1*⁺ LCMs at day 3, day 5, and day 8 following clodronate liposome treatment. Data are shown as mean ± s.e.m. (*N* = 3 independent experiments; *n* = 56 cells for day 3, *n* = 60 cells for day 5, and *n* = 60 cells for day 8). Statistical significance was determined by a two-tailed unpaired Student’s *t*-test; \*\**P* < 0.001, \*\*\**P* < 0.0001.

d. Quantification of cell circularity of RAW264.7 cells after long-term culture under different confinement conditions: control, slight confinement, moderate confinement, and strong confinement. Data are shown as mean ± s.e.m. (*N* = 3 independent experiments).

e. Schematic illustration of RAW264.7 cell migration under different confinement conditions: control, slight confinement, and strong confinement.

f. Mean instantaneous speed of RAW264.7 cells under control, slight confinement, moderate confinement, and strong confinement conditions. Data are shown as mean ± s.e.m. (*N* = 3 independent experiments).

g. Comparison of mean migration speed between non-transformed and transformed RAW264.7 cells under 3 μm strong confinement. Data are shown as mean ± s.e.m. (*N* = 3 independent experiments; *n* = 90 cells for non-transformed cells and *n* = 65 cells for transformed cells). Statistical significance was determined by a two-tailed unpaired Student’s *t*-test; \*\*\*\**P* < 0.0001.

h. Mean speed of RAW264.7 cells under different confinement conditions: control, slight confinement, moderate confinement, and strong confinement. Data are shown as mean ± s.e.m. (*N* = 3 independent experiments; *n* = 100 cells for Ctrl, *n* = 100 cells for Slight Conf., *n* = 65 cells for Moderate Conf., and *n* = 230 cells for Strong Conf.). Statistical significance was determined by a two-tailed unpaired Student’s *t*-test; ns, not significant; \*\*\**P* < 0.0001.

i. Representative images of THP-1 cells after long-term culture under different conditions: control, strong confinement, and PMA treatment.

j. Schematic illustration of cell morphology over 24 h under control (no confinement) and strong confinement (3 μm) conditions.

k. Quantification of cell circularity of THP-1 cells after long-term culture under control (no confinement) and strong confinement (3 μm) conditions. Data are shown as mean ± s.e.m. (*N* = 3 independent experiments).

Extended Data Figure 2. Supporting analyses of strong confinement induces macrophage-like activation in RAW264.7 cells.

a. Heat map showing the expression levels of macrophage activation – associated genes in the strong confinement versus control groups in RAW264.7 cells.

b. qPCR analysis of *Il1b*, *Tnf*, and *Nos2* expression in the control and strong confinement groups. Data are shown as mean ± s.e.m. (*N* = 3 independent experiments). Statistical significance was determined by a two-tailed unpaired Student’s t-test; *P < 0.01, ***P < 0.0001.

c. qPCR validation of *Nos2*, *Itgam*, and *Tnf* expression in the control, strong confinement, and LPS-treated groups. Data are shown as mean ± s.e.m. (*N* = 3 independent experiments). Statistical significance was determined by a two-tailed unpaired Student’s t-test; *P < 0.01, **P < 0.001, ***P < 0.0001.

Extended Data Figure 3. Long-term strong confinement regulates monocyte-to-macrophage differentiation in primary monocytes.

a,b. Representative images of Hoechst-labeled nuclear morphology in mMonocytes (a) and hMonocytes (b) after strong confinement. The red boxed regions show nuclei at the initial stage of confinement, adopting a characteristic horseshoe-shaped morphology. The blue boxed regions show nuclei after prolonged confinement, in which the horseshoe-shaped morphology is lost.

c. Representative images showing strong confinement–activated phagocytosis of cellular debris in hMonocytes.

d. Representative images of bone marrow–derived Lin⁻ cells, showing no detectable phagocytic activity under strong confinement.

e,f. Representative images of fluorescent bead phagocytosis (e) and quantification of phagocytic efficiency (f) in mMonocytes under control and strong confinement conditions. Data are shown as mean ± s.e.m. (*N* = 3 independent experiments). Statistical significance was determined by a two-tailed unpaired Student’s *t*-test; \*\*\*\**P* < 0.0001.

g. Quantification of fluorescence intensity of the macrophage marker F4/80 in mMonocytes and M-CSF – treated mMonocytes under control and strong confinement conditions. Data are shown as mean ± s.e.m. (*N* = 3 independent experiments; *n* > 40 cells for mMonocytes-Ctrl, *n* > 40 cells for mMonocytes-strong confinement, *n* > 40 cells for M-CSF-treated mMonocytes-Ctrl, and *n* > 40 cells for M-CSF-treated mMonocytes-strong confinement). Statistical significance was determined by a two-tailed unpaired Student’s *t*-test; \*\*\*\**P* < 0.0001.

h.i. Representative images (h) and quantitative analysis (i) of actin-based podosomes in mMonocytes, hMonocytes, and M-CSF–treated mMonocytes under control and strong confinement conditions. Data are shown as mean ± s.e.m. (*N* = 3 independent experiments; *n* = 42 cells for mMonocytes-Ctrl, *n* = 41 cells for mMonocytes-strong confinement, *n* = 47 cells for hMonocytes-Ctrl, *n* = 48 cells for hMonocytes-strong confinement, *n* = 43 cells for M-CSF-treated mMonocytes-Ctrl, and *n* = 41 cells for M-CSF-treated mMonocytes-strong confinement). Statistical significance was determined by a two-tailed unpaired Student’s *t*-test; \*\*\*\**P* < 0.0001.

Extended Data Figure 4. Supporting analyses of confinement-induced morphological and functional remodeling in hepatic-resident monocytes.

a. Giemsa staining illustrating the nuclear-to-cytoplasmic ratio of hepatic-resident monocytes (HRMs).

b. Immunofluorescence staining of HRMs showing *Cx3cr1* (green), CD11b (red), and nuclei labeled with DAPI (blue), reflecting their monocyte/macrophage identity.

c. Quantification of cell area of HRMs cultured under control (no confinement) or 3 μm strong confinement during long-term incubation. Data are shown as mean ± s.e.m. (*N* = 3 independent experiments; *n* = 56 cells for HRMs-0 h, *n* = 51 cells for HRMs-6 h, *n* = 51 cells for HRMs-12 h, *n* = 53 cells for HRMs-24 h; *n* = 54 cells for HRMs-strong confinement-0 h, *n* = 54 cells for HRMs-strong confinement-6 h, *n* = 50 cells for HRMs-strong confinement-12 h, and *n* = 52 cells for HRMs-strong confinement-24 h). Statistical significance was determined by a two-tailed unpaired Student’s *t*-test; ****P < 0.0001.

d. Representative images showing non-confind control have no phagocytosis of cellular debris in HRMs.

e. Representative images showing cell morphology after release from confinement following 48 h of strong confinement, illustrating partial recovery of cellular morphology.

Extended Data Figure 5. Supporting analyses of confinement-induced heterochromatin remodeling in monocytes.

a. Representative images of nuclear morphology in mMonocytes under control and strong confinement conditions.

b. Representative fluorescent images of H3K27me3 in M-CSF-treated mMonocytes under strong confinement. The dotted line marks the boundary between confined and non-confined regions.

c. Representative fluorescent images (left) and quantitative analysis (right) of H3K27me3 in RAW264.7 cells under control, slight confinement, and strong confinement conditions. Data are shown as mean ± s.e.m. (*N* = 3 independent experiments; *n* = 44 cells for Ctrl, *n* = 40 cells for slight confinement, *n* = 61 cells for strong confinement). Statistical significance was determined by a two-tailed unpaired Student’s *t*-test; ns, not significant; ****P < 0.0001.

d. Representative fluorescent images (left) and quantitative analysis (right) of H3K9me3 in RAW264.7 cells under control, slight confinement, and strong confinement conditions. Data are shown as mean ± s.e.m. (*N* = 3 independent experiments; *n* = 40 cells for Ctrl, *n* = 51 cells for slight confinement, *n* = 28 cells for strong confinement). Statistical significance was determined by a two-tailed unpaired Student’s *t*-test; ns, not significant.

e–h. Representative fluorescent images of CBX2 under strong confinement (e), CBX1 under strong confinement (f), CBX2 under non-confined control (g), and relative fluorescence intensity analysis (h) in RAW264.7 cells.

i,j. Representative fluorescent images (i) and relative fluorescence intensity analysis (j) of H3K27ac in RAW264.7 cells under control and strong confinement conditions. Data are shown as mean ± s.e.m. (*N* = 3 independent experiments; *n* = 68 cells for Ctrl and *n* = 86 cells for strong confinement). Statistical significance was determined by a two-tailed unpaired Student’s *t*-test; ns, not significant; ****P < 0.0001.

k. Heat map showing expression of heterochromatin-related genes in RAW264.7 cells under control and strong confinement conditions.

l. Relative mRNA expression levels of Ezh2, Suz12, Eed, Rbbp4, Rbbp7, Kdm6a, and Kdm6b in RAW264.7 cells under control and strong confinement conditions. Data are shown as mean ± s.e.m. (*N* = 3 independent experiments). Statistical significance was determined by a two-tailed unpaired Student’s *t*-test; ns, not significant; *P < 0.05, **P < 0.01, ***P < 0.001, ****P < 0.0001.

Extended Data Figure 6. Genetic and pharmacological modulation of Kdm6b during confinement-driven monocyte differentiation.

a. Representative fluorescent images of H3K27me3 in mMonocytes treated with DMSO or 10 μM GSKJ4 under non-confined control or strong confinement conditions.

b,c. Quantitative analysis of H3K27me3 fluorescence intensity in mMonocytes treated with DMSO or 10 μM GSKJ4 under non-confined control (b) and strong confinement (c) conditions. Data are shown as mean ± s.e.m. (*N* = 3 independent experiments). Statistical significance was determined by a two-tailed unpaired Student’s *t*-test; ****P < 0.0001.

d. Quantification of the proportion of cells exhibiting increased migration speed in RAW264.7 cells under strong confinement, comparing DMSO-treated control and 5 μM GSKJ4-treated conditions. Data are shown as mean ± s.e.m. (*N* > 3 independent experiments). Statistical significance was determined by a two-tailed unpaired Student’s *t*-test; ****P < 0.0001.

## Supplementary Video Legends

**Supplementary Video 1.** Representative time-lapse recording of RAW264.7 cell morphology under different confinement conditions: control, slight confinement, moderate confinement, and strong confinement, related to Fig. 1g.

**Supplementary Video 2.** Representative time-lapse recording of phagocytosis of cellular debris by RAW264.7 cells and THP-1 cells under strong confinement, related to Fig. 1j.

**Supplementary Video 3.** Representative time-lapse recordings of mMonocytes pretreated with 20 ng/mL M-CSF for 48 h prior to confinement, imaged under non-confined control and strong confinement conditions for an additional 48 h, related to Fig. 3b.

**Supplementary Video 4.** Representative time-lapse recordings showing the absence of cellular debris clearance by mMonocytes under control conditions and phagocytic clearance of cellular debris by mMonocytes under strong confinement, related to Fig. 3d.

**Supplementary Video 5.** Representative time-lapse recordings showing the absence of cellular debris clearance by hMonocytes under control conditions and phagocytic clearance of cellular debris by hMonocytes under strong confinement, related to Fig. S3c.

**Supplementary Video 6.** Representative time-lapse recordings showing accumulation of cellular debris due to cell death in mMonocytes under control conditions over 42 h, and phagocytic clearance of cellular debris by mMonocytes under strong confinement, related to Fig. 3e.

**Supplementary Video 7.** Representative time-lapse recordings showing dynamic changes in cell morphology of HRMs under control and strong confinement conditions, related to Fig. 4b.

**Supplementary Video 8.** Representative time-lapse recordings showing the absence of cellular debris clearance by HRMs under control conditions and phagocytic clearance of cellular debris by HRMs under strong confinement, related to Figs. 4d and S4d.

