## Supplementary figures and images for "Mechanical confinement drives monocyte-to-macrophage differentiation"

### Extended Data Figure 1

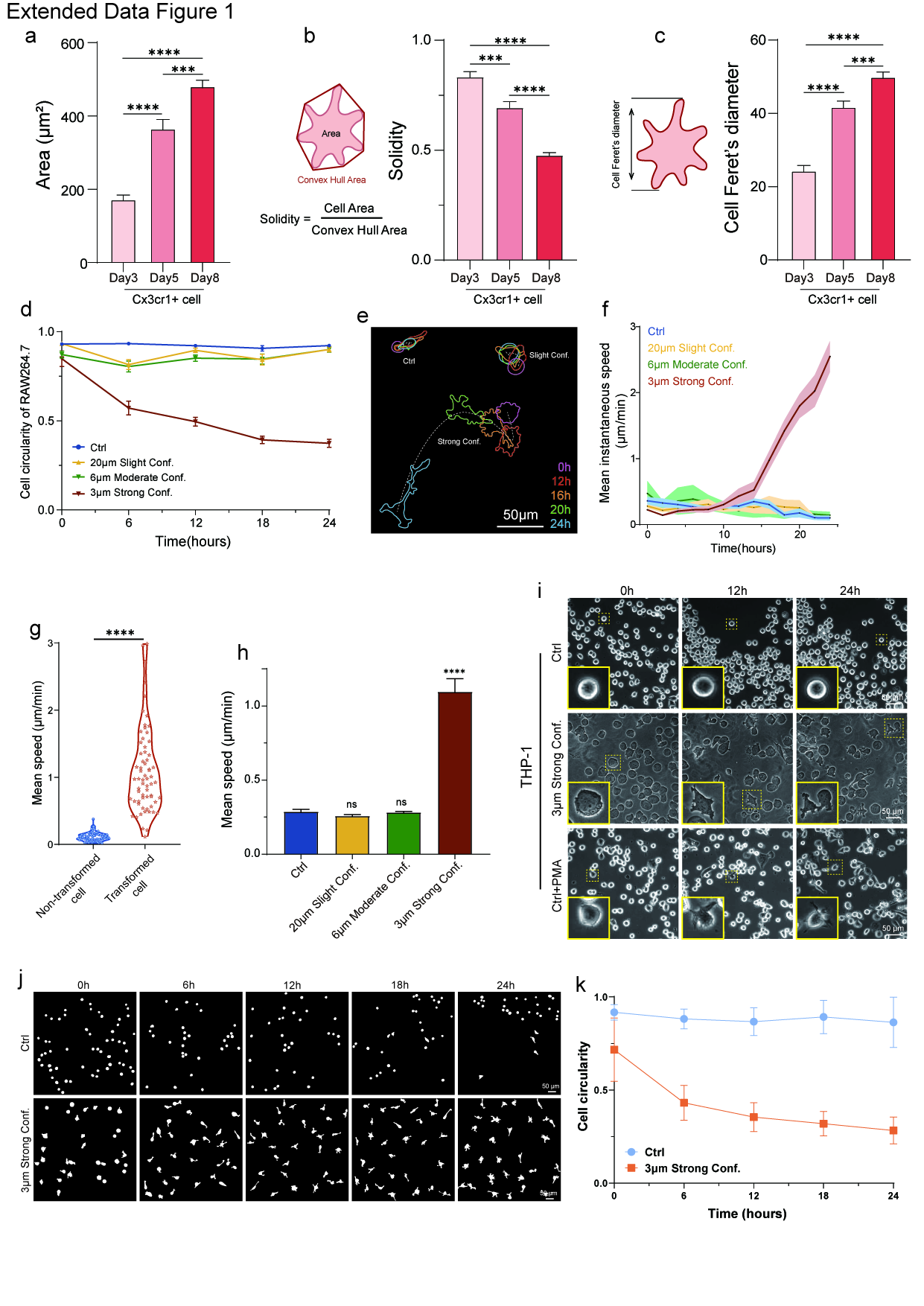

### Extended Data Figure 2

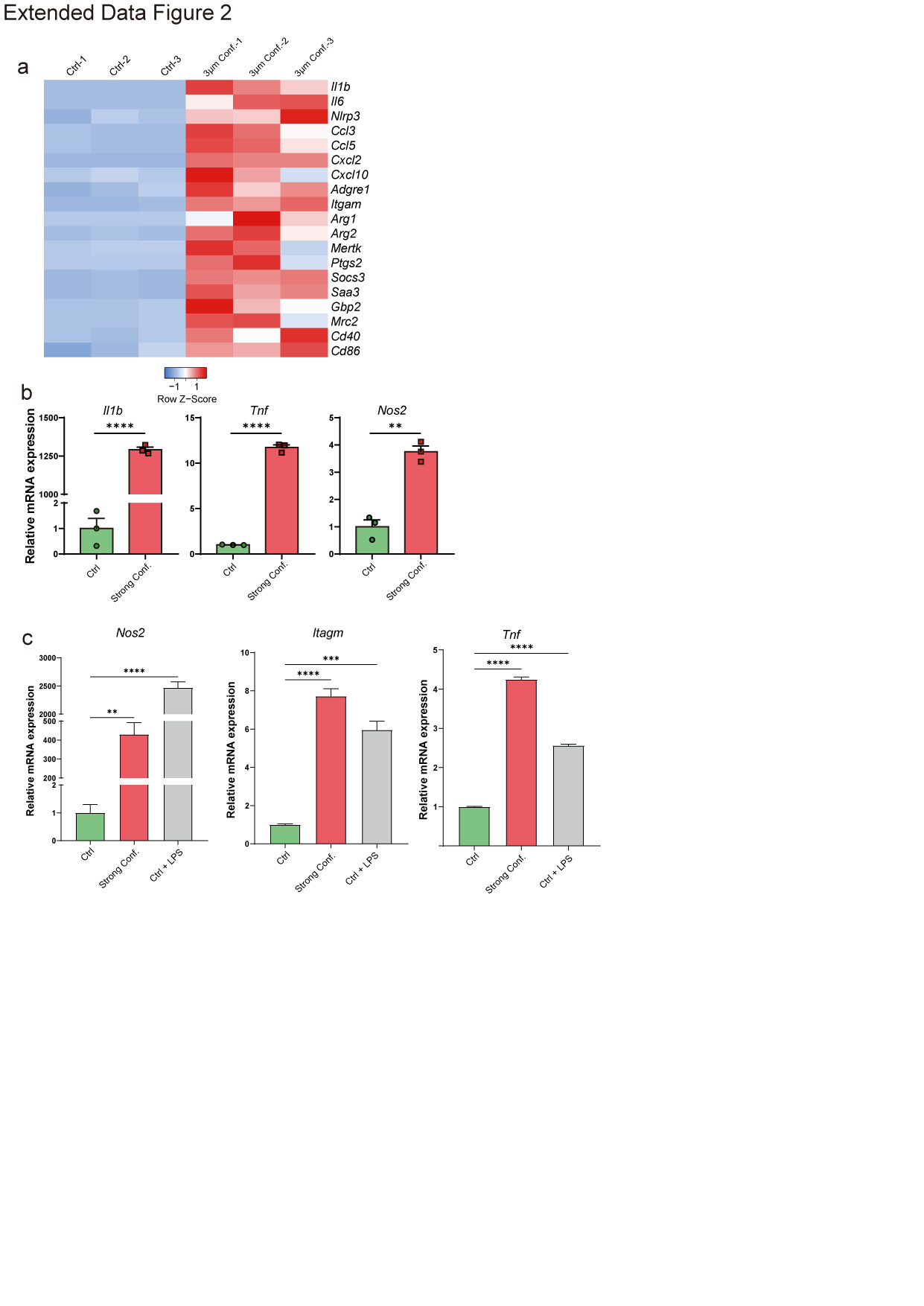

### Extended Data Figure 3

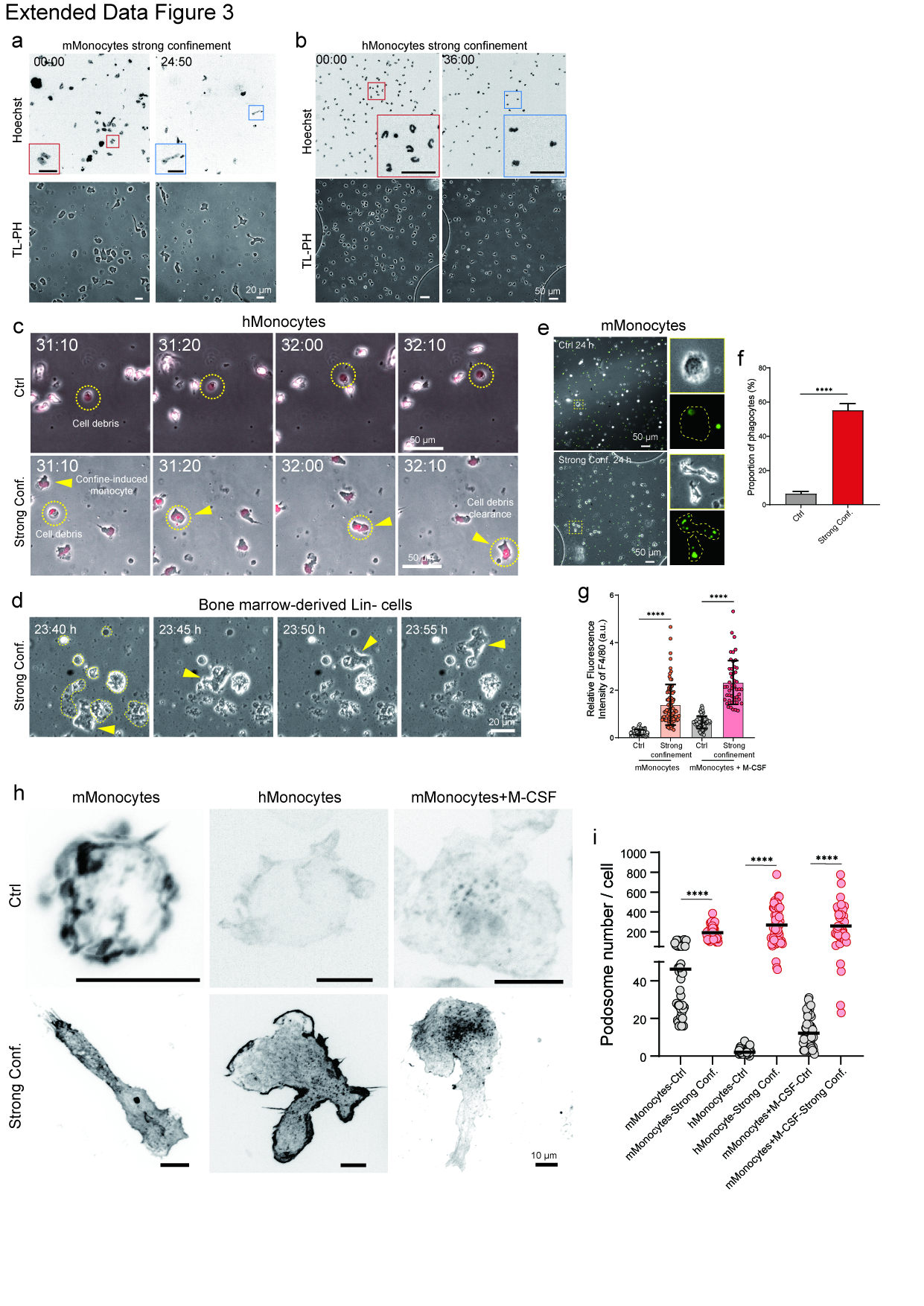

### Extended Data Figure 4

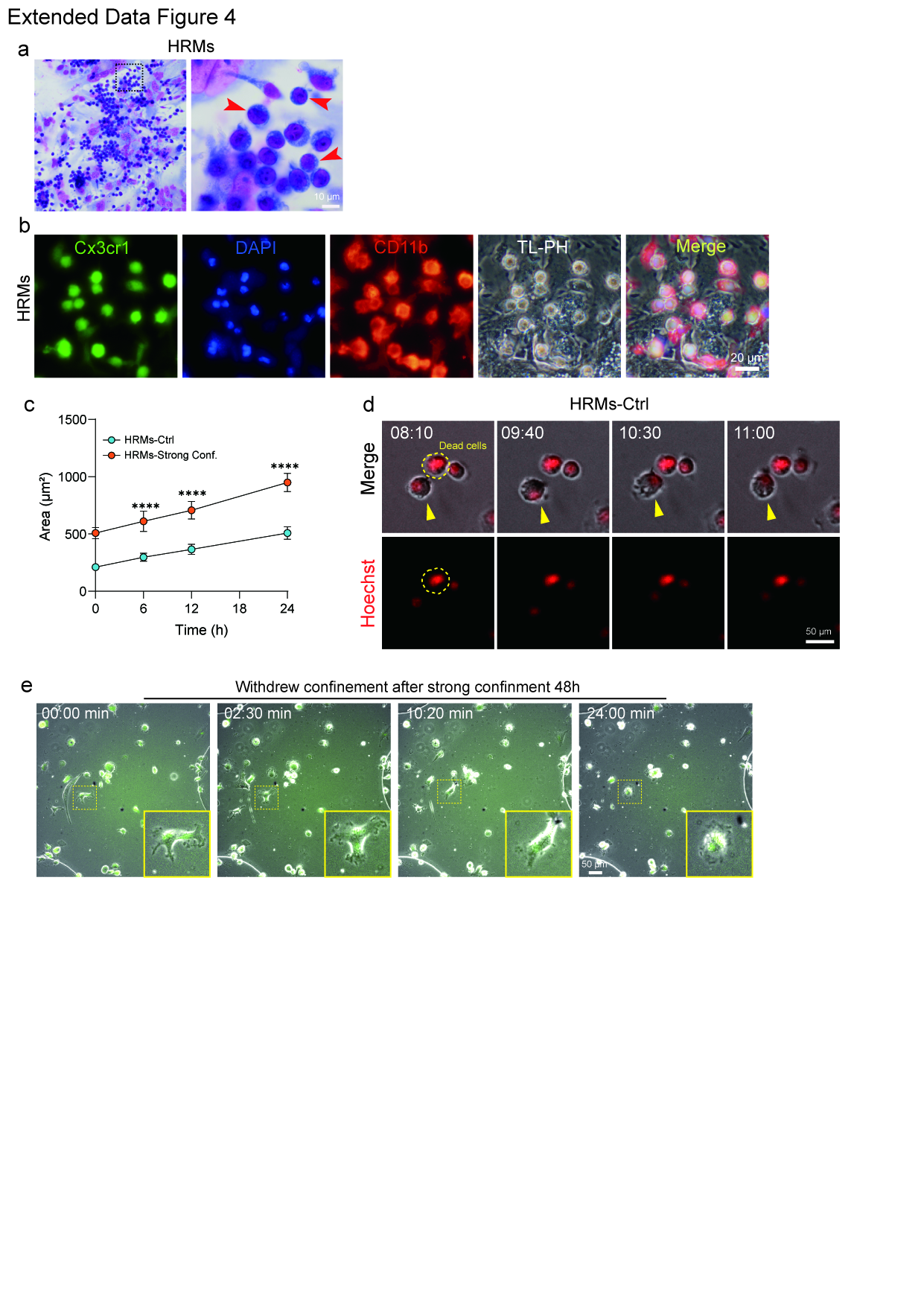

### Extended Data Figure 5

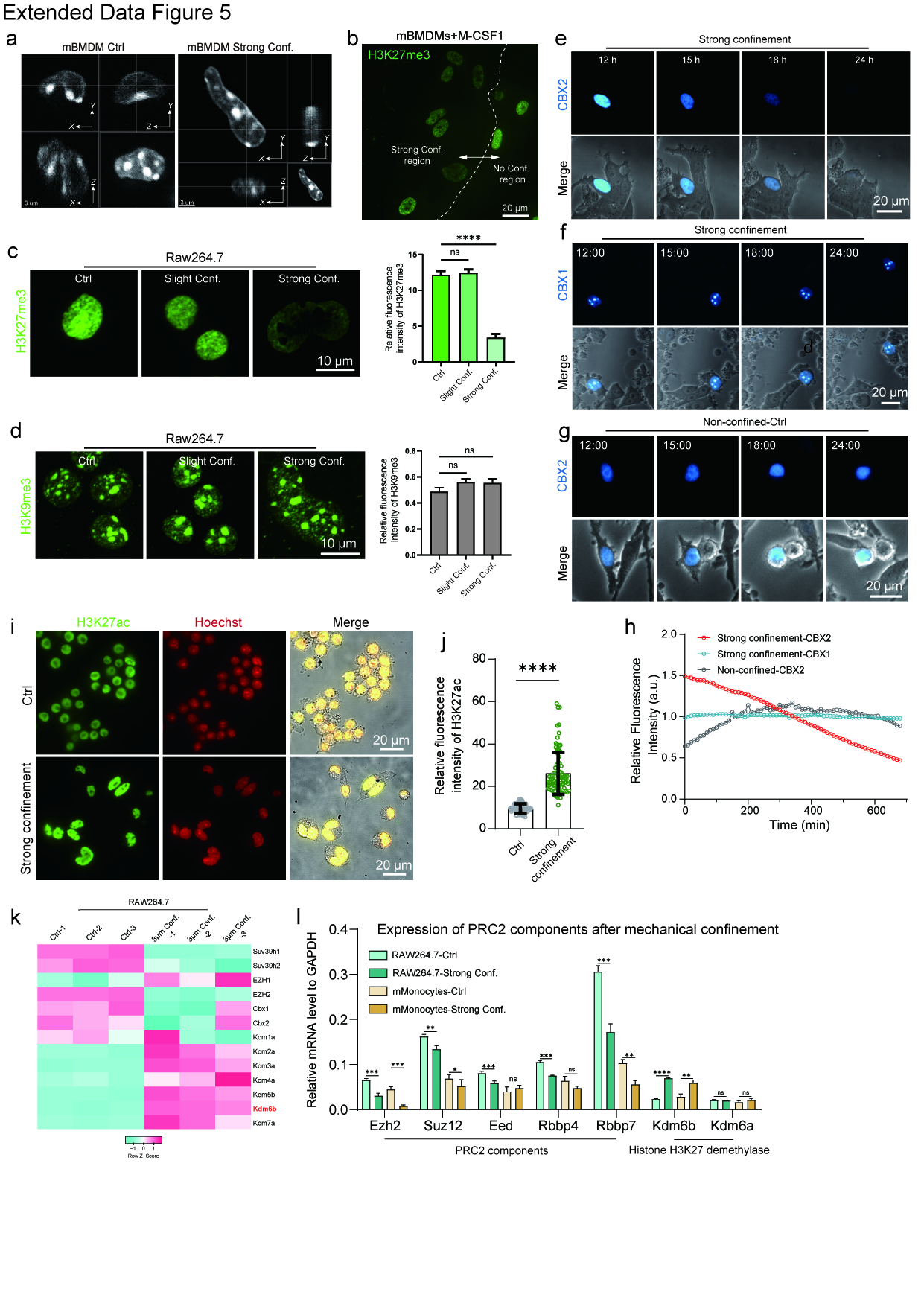

### Extended Data Figure 6

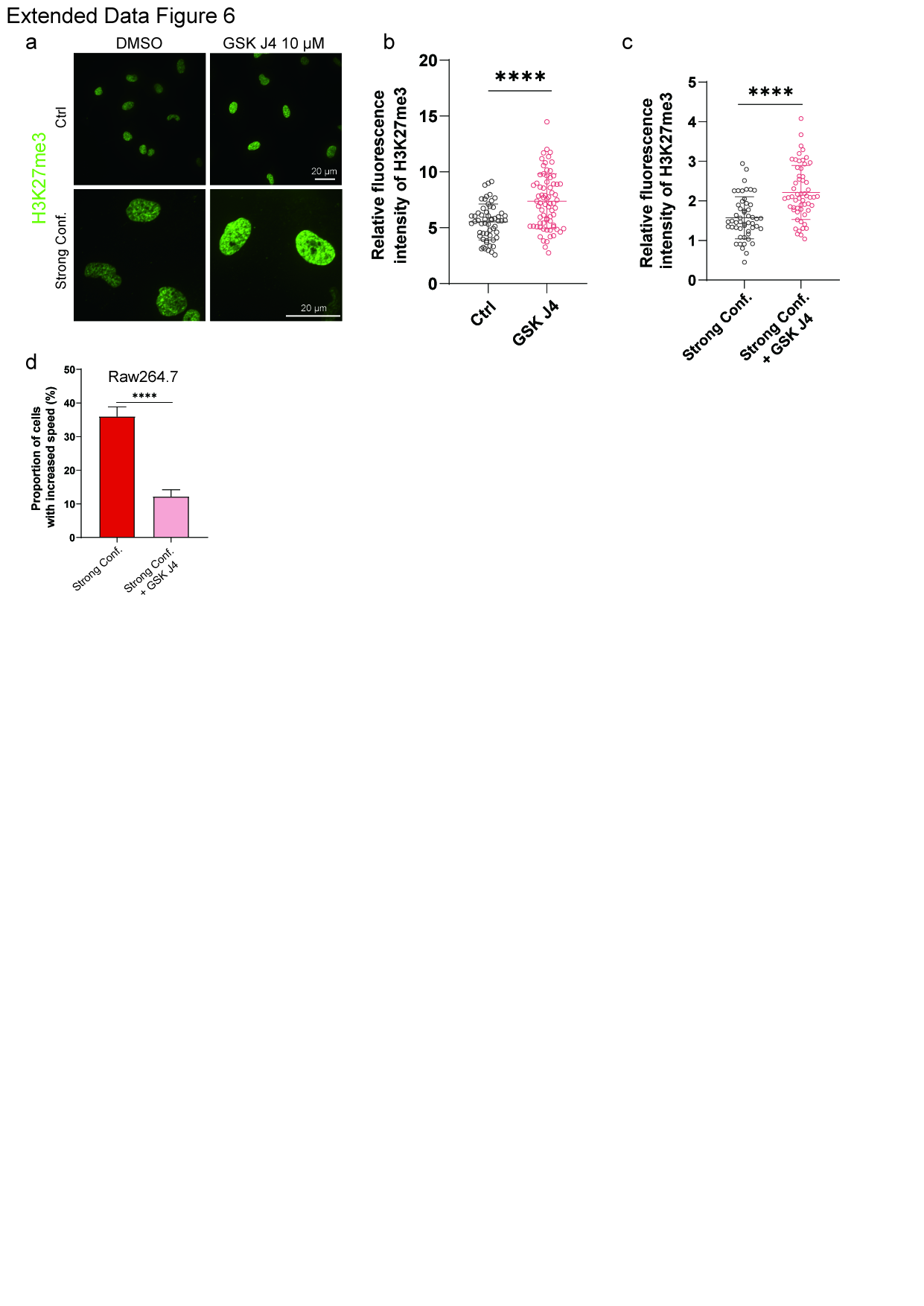
